# The zebrafish *dmrt* family genes have cooperative and antagonistic roles in sex determination and oogenesis

**DOI:** 10.1101/2022.08.28.505603

**Authors:** Jocelyn S. Steinfeld, Keith K. Ameyaw, Christopher G. Wood, Ryan M. Johnston, Ana J. Johnson Escauriza, Emma G. Torija, Kiloni Quiles, Kavita Venkataramani, Jessica N. MacNeil, Kellee R. Siegfried

**Author notes:** **Corresponding Author:** Kellee R. Siegfried, Biology Department, University of Massachusetts Boston Boston, MA 02125, USA.

## Abstract

The *double-sex and mab3 related transcription factor (dmrt)* gene family has conserved roles in sex determination and gonad development across metazoans. In zebrafish, *dmrt1* was previously shown to function in male sex-determination and testes development. To gain a broader knowledge of this gene family in sexual development, we investigated potential roles of all zebrafish *dmrt* family genes in sex-determination and gonad development using mutant analysis. The *dmrt2a* and *dmrt5* mutants conferred lethality prior to sex differentiation, whereas *dmrt2b* and *dmrt3* mutants were viable and fertile. *Dmrt2b* mutants had normal sex ratios while *dmrt3* showed slightly skewed sex ratios in some experiments, indicating that *dmrt3* has a minor role in sex-determination. We report a previously unknown role for *dmrt1* in ovary development. Although *dmrt1* mutant females were fertile, oogenesis did not progress normally, as evident from abnormal proportions of differently-staged oocytes within mutant ovaries. We also asked if *dmrt1* mutant phenotypes could be modified by loss of another *dmrt* family member. Analysis of *dmrt1;dmrt2a* mutants was possible as these double mutants were sub-viable, showing a partial rescue of the *dmrt2a* lethality in the *dmrt1* mutant background. The *dmrt1;dmrt2a* mutants had less severe female bias than *dmrt1* mutants suggesting that *dmrt2a* acts antagonistically to *dmrt1* in sex determination. Double mutants of *dmrt1* with either *dmrt2a* or *dmrt3* had more severe oogenesis defects than *dmrt1* mutants and had either sub-fertility with reduced fecundity or failed to breed, respectively. This study reveals previously unknown roles of zebrafish *dmrt1, dmrt2a*, and *dmrt3* in oogenesis.

## INTRODUCTION

The methods by which animals determine sex are simultaneously conserved and highly variable; while primary sex determination signals vary, many species demonstrate conservation in the genes acting downstream of this signal. The *dmrt (doublesex and mab2 related transcription factor)* family of genes is conserved throughout bilateria. Multiple members of the *dmrt* family have been implicated in sexual development in vertebrates, which comprises the developmental events that lead to sex determination and sexual differentiation (Picard *et al*. 2015). The defining feature of the Dmrt protein family is a conserved DNA binding domain, called the DM domain. Despite conservation of the DM domain, Dmrt proteins vary extensively in length and sequence (Mawaribuchi *et al*. 2019). This conservation results in the ability of Dmrt proteins to bind similar consensus binding sites, even between species (Murphy *et al*. 2007). Dmrt proteins often bind the promoters of both themselves and other Dmrt coding genes, suggesting auto- and cross-regulation occurs commonly within this family (Murphy *et al*. 2010). This suggests that Dmrt proteins have conserved functions as transcriptional regulators both within the gene family and across species. Differences in their function are likely derived through regulatory control, location of binding sites within the genome, and protein partners.

Functional interrogation of the *dmrt* family has been somewhat of a patchwork, with some genes thoroughly interrogated while the function of others remains poorly understood. The most well-studied member of this family is *dmrt1*, which has conserved roles in specifying and maintaining male cell fates in animals, particularly in the testis (Matson and Zarkower 2012; Picard *et al*. 2015). The *Dmrt6* and *Dmrt7* genes also have documented roles in testis development, where they are important for mammalian spermatogenesis (Kim *et al*. 2007a; Zhang *et al*. 2014a, 2014b; Li *et al*. 2020). Additional *dmrt* genes are implicated in ovary development, though their roles are not fully understood. For example, loss of *Dmrt4* in mice produces polyovular follicles in ovaries and human *DMRT5, DMRT6*, and *DMRT7* have been suggested to be involved in female germ cell development (Balciuniene *et al*. 2006; Poulain *et al*. 2014). However, outside of mammals, the function of most *dmrt* genes in sexual development remains unknown. Of the eight *dmrt* genes found in vertebrates, zebrafish have five members of this family: *dmrt1*, *dmrt2a*, *dmrt2b*, *dmrt3*, and *dmrt5* (Figure 1)(Huang *et al*. 2002). An in-depth functional analysis of this group of genes will aid our understanding of this highly conserved gene family.

**Figure 1.**
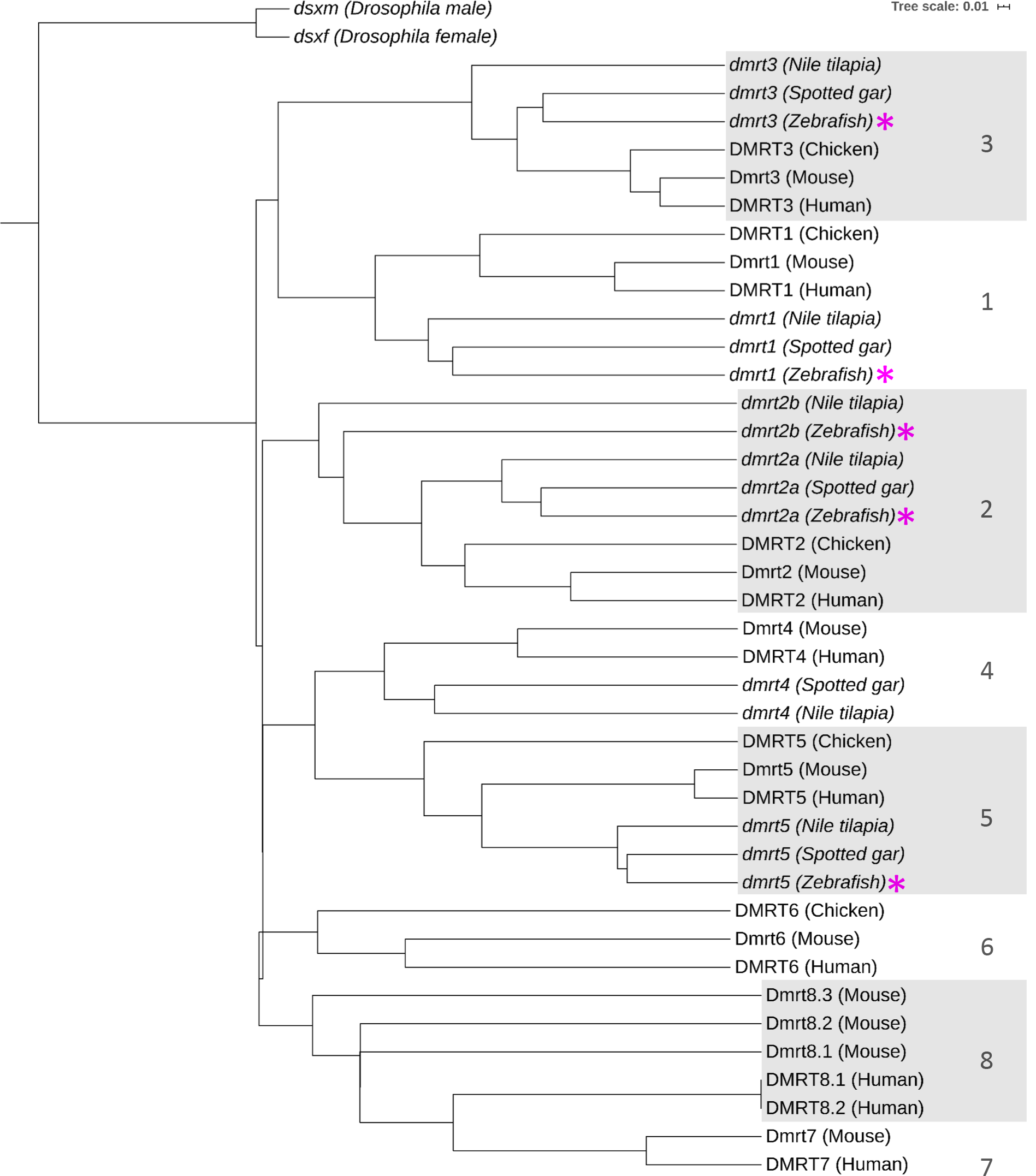
Zebrafish have five *dmrt* genes. Zebrafish *dmrt* genes cluster with their orthologues in other vertebrate species. Pink asterisk = zebrafish gene. Scale bar of 0.01 = 1% sequence difference

Zebrafish *dmrt1* has a role in male sexual development, similar to other vertebrate animals. In mice, *Dmrt1* is required for the maintenance of male cell fates and for germ cell survival and sperm differentiation (Raymond *et al*. 2000; Kim *et al*. 2007a; Matson *et al*. 2010, 2011; Takashima *et al*. 2013), Interestingly, loss of *Dmrt1* in testicular Sertoli cells of adult mice causes these cells to transdifferentiate to an ovarian granulosa cell fate, indicating that it is required throughout the animal’s life to maintain testis versus ovary cell fates (Matson *et al*. 2011). *Dmrt1* has minor roles in ovary development, however mutations disrupting this gene do not affect female fertility (Krentz *et al*. 2013). Similar to mice, loss of *Dmrt1* in humans results in testicular dysgenesis, suggesting the function of *Dmrt1* is conserved in mammals (Veitia *et al*. 1997; Tannour-Louet *et al*. 2010; Murphy *et al*. 2015). In birds, *DMRT1* is the primary testis-determining gene, where two copies on the avian sex-linked Z chromosome are required to specify testis fate (Smith *et al*. 2009). Interestingly, it is also required for early formation of the chicken Müllerian ducts (Ayers *et al*. 2015). This finding suggests that, as in mice, avian *DMRT1* is important for development in both sexes. The zebrafish *dmrt1* gene functions in male fate determination and testis development (Webster *et al*. 2017; Lin *et al*. 2017). The majority of *dmrt1-/-* develop as fertile females while a small percentage of mutants are sterile males. These males exhibit testicular dysgenesis and complete loss of sperm. Despite having a critical role in male development, some *dmrt1-/-* zebrafish still develop as sterile males, demonstrating that male fate can be specified in the absence of *dmrt1*.

Two paralogues of *dmrt2* are present in zebrafish, *dmrt2a* and *dmrt2b.* This duplication occurred independently and prior to the teleost genome duplication event (Zhou *et al*. 2008; Johnsen and Andersen 2012). Morpholino knockdowns have implicated *dmrt2a* in somitogenesis, left-right asymmetry during segmentation, and fast muscle differentiation (Meng *et al*. 1999; Saúde *et al*. 2005; Lu *et al*. 2017). Zebrafish *dmrt2b* has been suggested to be involved in hedgehog signaling, left-right asymmetry, and cranial-facial development (Liu *et al*. 2009; Li *et al*. 2018). In mice, *Dmrt2* is required for embryonic myogenesis and in humans and mice it is necessary for vertebrae and rib development (Sato *et al*. 2010; Bouman *et al*. 2018). In the mouse gonad, *Dmrt2* exhibits strong testicular expression with weaker expression in the ovary (Kim *et al*. 2003). Conversely, data suggests that human DMRT2 is expressed exclusively in the fetal ovary during the first and second trimester (Poulain *et al*. 2014). Zebrafish *dmrt2a* and *dmrt2b* are both expressed in the gonads, but it is unclear what role they may play in sexual development (Liu *et al*. 2009).

In several animals, *dmrt3* functions in brain development. In the mammalian brain, it is required for normal development of the telencephalon in mice, and normal locomotion in mice and horses (Andersson *et al*. 2012; Konno *et al*. 2012; Saulnier *et al*. 2013; De Clercq *et al*. 2018; Desmaris *et al*. 2018; Perry *et al*. 2019). This function is somewhat conserved in zebrafish, where *dmrt3* is important for larval fish locomotion and juvenile swim performance, though zebrafish lose brain expression of *dmrt3* by adulthood (Li *et al*. 2008; Del Pozo *et al*. 2020). Interestingly, murine *Dmrt3* is expressed in the developing ovaries and testes, but expression is lost at maturity (Kim *et al*. 2003). Conversely, human DMRT3 is expressed in ovaries but not testes, at least through the first two trimesters (Poulain *et al*. 2014). *Dmrt3* null mutant mice demonstrated no gonadal defects, and mice mutant for both *Dmrt1* and *Dmrt3* had no modification of the *Dmrt1* phenotype (Inui *et al*. 2017). Zebrafish *dmrt3* is expressed in both ovaries and testes, however its function in sexual development remains unclear (Li *et al*. 2008).

*Dmrt5* also has known roles in the brain. Mammalian *Dmrt5* is expressed in the brain and functions cooperatively and critically with *Dmrt3* in hippocampal, neuronal, and cerebral cortex development (Konno *et al*. 2012; Saulnier *et al*. 2013; Muralidharan *et al*. 2017; De Clercq *et al*. 2018; Desmaris *et al*. 2018; Ratié *et al*. 2020). *Dmrt5* mutant mice exhibited almost complete lethality within a few hours of birth due to brain defects (Saulnier *et al*. 2013). The function of *dmrt5* in brain development appears conserved in zebrafish, where it is required for neurogenesis, as well as proper specification of corticotropes and gonadotropes. Similar to mice, zebrafish *dmrt5* mutants were larval lethal with defects in telencephalon and pituitary development, failures in corticotrope differentiation, and reduced expression of telencephalic neural markers (Yoshizawa *et al*. 2011; Graf *et al*. 2015). Zebrafish *dmrt5* is expressed in germ cells of adult ovaries and testes and has a role in regulating gene expression in the testis (Guo *et al*. 2004; Xu *et al*. 2013).

In this study, we aimed to determine effects of loss of *dmrt2a, dmrt2b, dmrt3*, and *dmrt5* in zebrafish sex determination and gonad differentiation by characterizing mutations disrupting these genes. Furthermore, we sought to determine if *dmrt1* acts redundantly with other *dmrt* genes in the zebrafish gonad by generating double mutants with *dmrt1*. Here, we report that mutations disrupting *dmrt2a* and *dmrt5* are larval lethal between 7 and 12 days post fertilization (dpf), precluding analysis of sexual development. Mutations affecting *dmrt2b* and *dmrt3* were adult viable with little to no apparent defects in sexual development. Double mutant analysis with *dmrt1* revealed that *dmrt1*, *dmrt2a*, and *dmrt3* have roles in ovary development. In particular, the *dmrt2a-/-* lethality was suppressed in the *dmrt1* mutant line and these double mutants exhibited ovary defects. *Dmrt2a-/-* partially suppressed the female sex bias of *dmrt1-/-* and enhanced minor *dmrt1-/-* ovary defects, leading to impaired fertility and fecundity of double mutants. These data suggest that *dmrt2a* and *dmrt1* act antagonistically to each other in sex determination but synergistically in oogenesis. We also report that *dmrt1-/-;dmrt3-/-* exhibited more severe ovary defects than in *dmrt1-/-*, suggesting a role of *dmrt3* in ovary development as well. By contrast, neither *dmrt2a-/-* nor *dmrt3-/-* modified the *dmrt1-/-* testis defects. Through double mutant analysis with *dmrt1-/-*, we have revealed previously unidentified functions for *dmrt1, dmrt2a* and *dmrt3* in female development.

## METHODS

### Zebrafish lines

Zebrafish were raised and maintained in standard conditions. Institutional Animal Care and Use Committee approval was attained for all animal procedures. All experiments were performed in the wild-type AB line and mutant lines *dmrt1^uc27^ dmrt5^sa13156^*(Kettleborough *et al*. 2013; Webster *et al*. 2017). New mutations generated in this study were: *dmrt2a^umb1^, dmrt2a^umb11^, dmrt2a^umb12^, dmrt2b^umb4^, dmrt2b^umb6^, dmrt3^umb2^, dmrt3^umb3^, dmrt3^umb14^.* A complete table of primers is presented in Table S1.

The *dmrt1^uc27^* line was maintained as previously described (Webster *et al*. 2017). The *dmrt5^sa13156^* allele was provided by the Zebrafish International Resource Center (ZIRC) and was generated using ENU mutagenesis by the Zebrafish Mutation Project (Kettleborough *et al*. 2013). Fish harboring this allele were genotyped by amplification using Derived Cleaved Amplified Polymorphic Sequences (dCAPS) primers (Neff *et al*. 2002) JS42/JS43, and treated with BamHI-HF (NEB R3136S), which digests the wild-type amplicon. Products were visualized on a 7% polyacrylamide DNA gel.

The remaining mutations described in this study were generated in the AB line using CRISPR/Cas9 genome editing. All guide RNAs were designed downstream of potential alternative start codons that might result in production of functional proteins (i.e. ATG codons 5’ of the DM domain). To generate the *dmrt2a^umb1^* allele, a CRISPR guide RNA (gRNA) was designed to target the first exon of *dmrt2a* (5’- GGTGTTCACCGGCCCCGGAG-3’), and the gRNA was synthesized as described previously (Gagnon *et al*. 2014). Cas9 mRNA was generated using the plasmid *pCS2-nCas9n* (Addgene plasmid # 47929)(Jao *et al*. 2013) and mRNA was synthesized using the mMESSAGE mMACHINE™ SP6 Transcription Kit (Invitrogen AM1340). The CRISPR gRNA (100pg per embryo) and Cas9 mRNA (600pg per embryo) were injected into single-cell embryos. Three days post-injection, larvae were screened in pools using PCR with primers dmrt2a-1F/dmrt2a-1R, then annealed and assayed with T7 endonuclease (NEB M0302S) to look for mismatched nucleotide bases. Samples with multiple bands on gels were sub-cloned using CloneJET PCR Cloning Kit (Thermo Scientific K1231), and subsequently Sanger-sequenced to determine mutations. Genotyping for maintenance of the *dmrt2a^umb1^* line was done using primers JS36/JS37 or JS137/JS138 and treating the amplicon with enzyme BslI (NEB R0555S) which digests only the wild-type amplicon. Products were run on a 7% polyacrylamide DNA gel to visualize.

The *dmrt2a^umb11^, dmrt2a^umb12^, dmrt2b^umb4^*, *dmrt2b^umb6^*, *dmrt3^umb2^* and *dmrt3^umb3^* alleles were also generated using CRISPR/Cas9 mutagenesis in AB fish. The above-described target sequence for *dmrt2a* was used, and targets were selected for both *dmrt2b* (5’- CGGACGGGAGGTCATGGGCA -3’) and *dmrt3* (5’- GTGCGCGCTGCAGGAACCAC - 3’) in the first exons. One CRISPR crRNA for each gene was purchased from Integrated DNA Technologies (IDT). These crRNA were annealed to the IDT universal trcrRNA (IDT 1072532) per IDT protocol to form the gRNA complex. The CRISPR gRNA (5pg per embryo) and Cas9 mRNA (150pg per embryo, System Biosciences CAS500A-1) were injected into single-cell embryos. Injected embryos were screened individually at 1 day post-injection using PCR with primers JS36/JS37 for *dmrt2a*, JS54/JS62 for *dmrt2b* and JS30/JS31 for *dmrt3*. Samples were run on a 7% DNA polyacrylamide gel to visualize, and samples with more than one band were sub-cloned and sequenced as described above.

Genotyping for mutant lines was largely performed with the same PCR method as screening for mutations, with a few exceptions. For genotyping *dmrt2a^umb11^* and *dmrt2a^umb12^*, primers used were JS137/JS138, along with enzymatic digestion of the wild-type amplicon with enzyme BslI (NEB R0555S). Genotyping *dmrt2b^umb4^*required treatment with the enzyme BaeGI (NEB R0708S), which digests only the wild-type amplicon.

Double mutants for *dmrt1^uc27^*with *dmrt2a* and *dmrt3* were also generated with CRISPR/Cas9 mutagenesis. The same genomic targets, IDT CRISPR crRNA, and manually synthesized gRNA in the case of *dmrt2a*, were used to generate mutant alleles. *Dmrt1^uc27^* mutant females were crossed to *dmrt1^uc27/+^*males, and the resulting single-cell embryos were injected and screened as described above, with the addition of also genotyping for *dmrt1^uc27^*. Linked mutations were confirmed through identification of *dmrt1^uc27^* homozygous mutant females carrying novel mutations for *dmrt2a* or *dmrt3*, or by breeding *dmrt1^uc27^* heterozygous male carriers to homozygous females and genotyping embryos to confirm presence of homozygous *dmrt1* mutant progeny carrying the novel allele. Genotyping for maintenance of lines was performed with primers JS137/JS138 and enzyme BslI (NEB R0555S) for *dmrt2a^umb13^*, and JS30/JS31 for *dmrt3^umb14^* with enzyme BSAJI (NEB R0536S), which digests only the wild-type amplicon.

### Germ cell ablation

A morpholino targeting the *deadend (dnd)* transcript was injected into wild-type, single-cell embryos, as previously described (Siegfried and Nüsslein-Volhard 2008).

### Fertility assays

Adult fish were paired in individual mating boxes, with plastic plants. The following morning, eggs and embryos were collected, counted, and sorted by fertilization state, which is identifiable by 4-6 hours post-fertilization (hpf). Eggs that were cloudy and degrading or not developing through proper cleavage stages were considered unfertilized. Only clutches of 80 or more eggs/embryos were included, and percent fertilized is the number of live, developing embryos out of the total eggs/embryos counted per clutch. In the case of *dmrt1;dmrt2a* double mutants, abnormal/nonviable progeny were determined by counting again at 24 hpf, when various forms of abnormal development were readily evident.

### Histology and ovary quantification

Histology was performed on fish euthanized with tricaine (MS-222). Fish trunks were isolated, and abdomens were grossly examined to confirm sex. Samples were fixed either for 2-4 hours in Davidson’s solution (isolated gonads) or overnight in Bouin’s fixative (whole torsos) at room temperature, then embedded in paraffin. Testes were cut into 5µm sections and ovaries into 7 µm sections and were stained using Modified Harris hematoxylin and Eosin Y (H&E) (Siegfried and Steinfeld 2021).

Ovary quantification was performed on H&E sections. Three 7µm sections were taken from each sample, no less than 90µm apart to ensure no overlapping cells between sections. Slides were imaged on a Zeiss Axio-Observer Z1 inverted microscope with ZEN microscope software using a tile-scan approach, to produce an image of the entire section. Each germ cell was categorized by stage according to Selman *et al*. (Selman *et al*. 1993). The proportion of each oocyte stage out of the total number of germ cells in that sample was calculated and averaged across all samples for each genotype. Chi-square tests were performed to compare the overall changes in proportions between genotypes, and students’ T-tests were performed to determine significance between genotypes for each individual stage.

### RT-PCR

RNA was collected from whole organs. Ovaries were collected individually, and all other tissues were pools from multiple fish. RNA was isolated using TRI Reagent (Sigma 93289) according to the manufacturer’s protocol. Synthesis of cDNA was done through reverse-transcription using AMV Reverse Transcriptase (NEB M0277S). Analysis of qRT-PCR data was performed using GraphPad Prism software.

### qRT-PCR

RNA was collected from whole ovaries, except in the case of *dmrt1;dmrt2a* female fish, where only half of each ovary was collected. Ovaries and testes were collected individually. RNA was extracted using TRI Reagent (Sigma 93289) according to the manufacturer’s protocol. Total RNA was DNase treated using the TURBO DNA-free™ Kit (Invitrogen AM1907), and reverse-transcription cDNA synthesis was performed using the SuperScript™ IV Reverse Transcriptase (Invitrogen 18090010). qRT-PCR was performed using PowerUp™ SYBR™ Green Master Mix (Applied Biosystems A25742) and QuantStudio™ 3 Real-Time PCR System (Applied Biosystems A28567). All primers used in qRT-PCR had primer efficiencies above 98%, calculated using standard ΔΔCT methods.

### Phylogenetic analysis

The phylogeny was constructed using protein sequences retrieved from Ensembl (https://useast.ensembl.org/). All relevant ensemble transcript IDs are contained in Table S2. Phylogeny construction was performed using the phyloTv2 tree generator (https://phylot.biobyte.de/).

## RESULTS

### Expression of dmrt genes in adult zebrafish

Zebrafish have five *dmrt* family genes, all of which have orthologues in other vertebrates (Figure 1). Given that *dmrt1* frequently plays a role in sex determination and gonad development across metazoans, including in zebrafish, and that other *dmrt* genes have known roles in the gonad in some animals, we asked if the other four members of the zebrafish *dmrt* family (*dmrt2a, dmrt2b, dmrt3*, and *dmrt5)* also play a role in gonadal sex determination or development. Given that these genes have yet to be analyzed simultaneously in a single study, we first tested if these genes are expressed in the zebrafish gonad (Figure 2). By RT-PCR on tissues from wild-type adults, we found that all five zebrafish *dmrt* genes were expressed in both adult ovary and testis, suggesting that they may function in the gonad. Previously, *dmrt1* and *dmrt5* were shown to be expressed in germ cells of adult gonad (Guo *et al*. 2005; Yoshizawa *et al*. 2011; Xu *et al*. 2013; Webster *et al*. 2017). We detected expression of all genes in testes lacking the germline, demonstrating that these genes are expressed in the somatic cells of male gonads (Figure 2). Because zebrafish gonads lacking germ cells always develop as testes, we could not generate ovaries lacking germ cells (Siegfried and Nüsslein-Volhard 2008). These genes may be expressed in other cell types in the gonad as specific expression in other cell types of the gonads was not tested. We also tested a panel of additional adult organs to determine the expression of these genes outside the gonad. We found that while *dmrt2a, dmrt2b*, and *dmrt5* are expressed in numerous tissues, *dmrt1* and *dmrt3* were expressed only in the gonad (Figure 2). To determine if these genes were expressed at different levels between ovaries and testes, we analyzed previously reported RNA-sequencing data on isolated adult zebrafish ovaries and testes (Yan *et al*. 2019). *Dmrt1* was strongly enriched in testes compared to ovaries, as previously reported, while other *dmrt* genes had lower expression levels in both sexes (Figure 2) (Guo *et al*. 2005; Webster *et al*. 2017). *Dmrt2a* exhibited slightly higher expression in testes than ovaries. *Dmrt2b* was expressed in ovaries but not detected in testes, whereas *dmrt3* was enriched in testes and not detected in ovaries. *Dmrt5* was expressed at low levels in both sexes. We conclude that all the *dmrt* genes are expressed in the gonad with some genes enriched in one sex. These data suggest that each of these genes have the potential to play roles in sex determination and gonad development.

**Figure 2.**
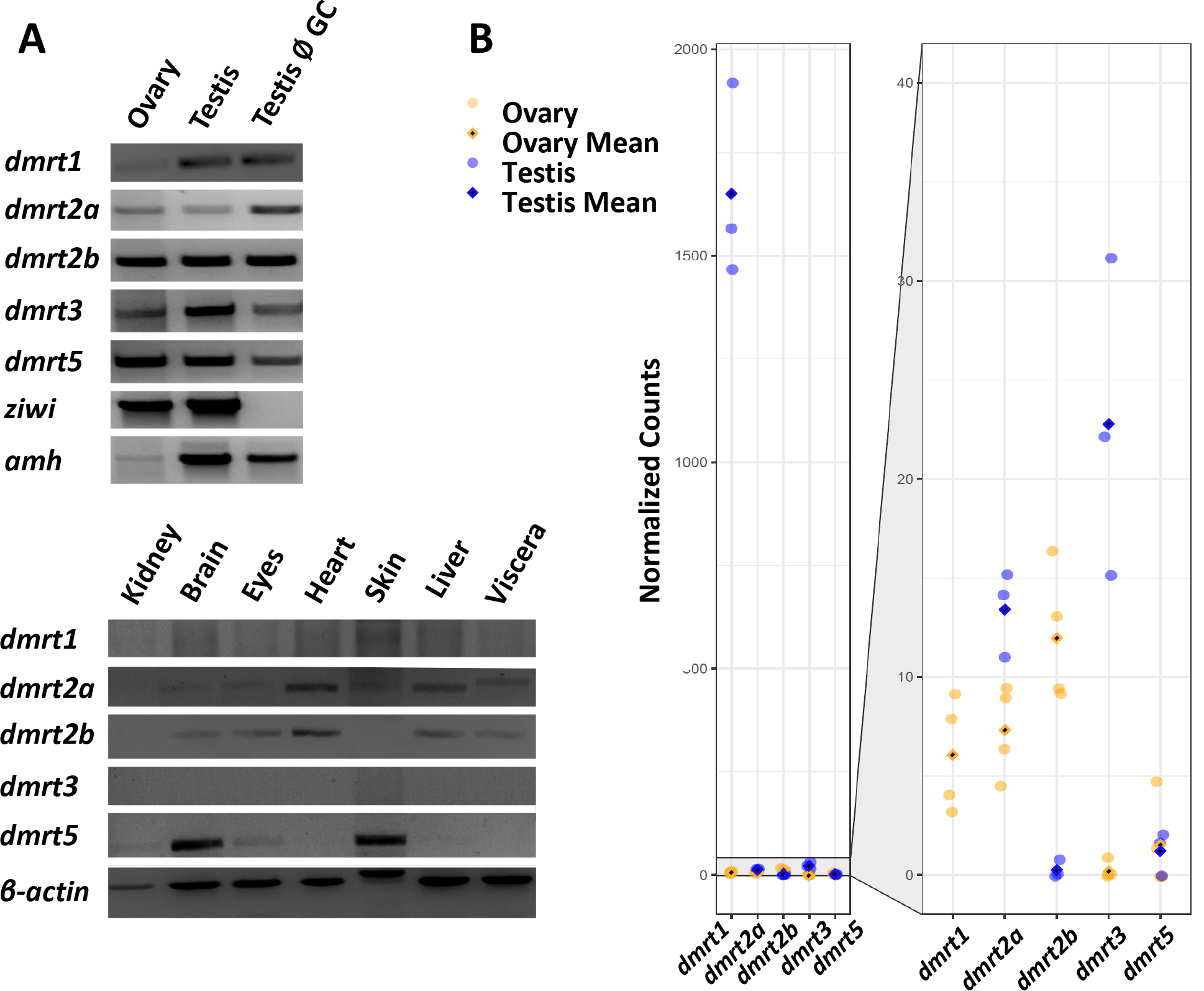
All zebrafish *dmrt* genes are expressed in the gonads and have varying expression profiles in other tissues. A) RT-PCR results showing the expression of the *dmrt* genes in various tissues. *β-actin* is a ubiquitously expressed gene; *ziwi* is a marker for germ cells; *amh* is a marker for testis Sertoli cells and ovarian follicle cells, testis Ø GC= testis with ablated germline. B) Analysis of RNAseq data from adult testes and ovary. *Dmrt1* has the highest expression levels in testes and lower in ovaries. Other *dmrt* genes are less robustly expressed in gonads. *Dmrt2b* has higher expression in ovaries while *dmrt3* is more abundant in testes than ovaries. *Dmrt2a* and *dmrt5* are expressed at similar levels in both sexes.

### Dmrt2a and dmrt5 mutants are homozygous larval lethal

Mutants disrupting each zebrafish *dmrt* family gene were generated or acquired in order to assess gene function. The null allele *dmrt1^uc27^* has been previously characterized by our lab, and the predicted-null allele *dmrt5^sa13156^*was produced by the Zebrafish Mutation Project and obtained from the Zebrafish International Resource Center (ZIRC) (Kettleborough *et al*. 2013). We generated two mutant alleles each for *dmrt2a*, *dmrt2b*, and *dmrt3* using CRISPR/Cas9 genome editing, and all are predicted to produce truncated proteins that disrupt function (Figure S1, Figure 3). To determine the necessity of *dmrt2a, dmrt2b, dmrt3*, and *dmrt5* in development, we first assessed viability of homozygous mutants. For homozygous viable mutations, we assayed family sex ratios, fertility, and gonad histology to ask if any of these genes had necessary functions in sexual development.

**Figure 3.**
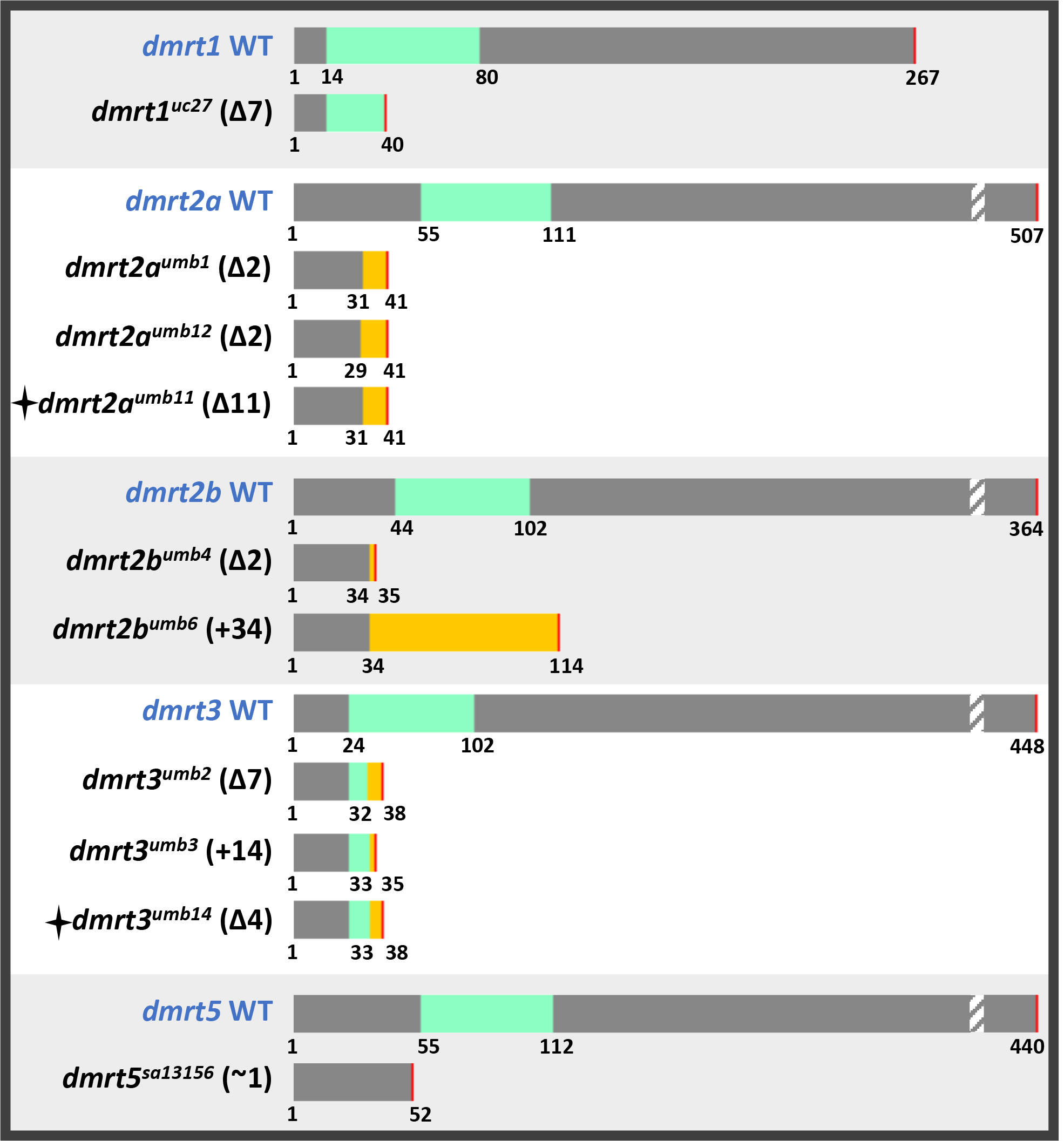
All *dmrt* mutants used in this study. All *dmrt* mutations result in truncated proteins and disruption of the DM domain. Grey = protein sequence, green = DM domain, yellow = frame-shifted protein sequence, red = stop, star = linked mutation with *dmrt1.* Numbers denote amino acid position. Drawings are roughly to scale, hatched sections denotes that the full sequence is longer than shown.

We found that *dmrt2a* and *dmrt5* mutants were larval lethal, whereas *dmrt2b* and *dmrt3* were adult viable. To assess viability, we analyzed genotypic ratios of adult fish derived from incrossing heterozygous parents. Both *dmrt2b* mutants and one *dmrt3^umb3^* mutant were adult viable and present at expected frequencies, however the *dmrt3^umb2^* mutants were present at a lower-than-expected frequency (Table 1). To address this, we generated trans-heterozygous *dmrt3^umb2/umb3^* fish, and analyzed their genotypic ratios in comparison to heterozygous and wild-type siblings. Fish that carried the *dmrt3^umb2^* allele, either as homozygous mutants or as trans-heterozygotes with *dmrt3^umb3^*, were consistently underrepresented in the population, suggesting that the chromosome harboring this allele confers some lethality when a wild-type allele is absent (Table 1). Previously it was reported that *dmrt5^ha2^*mutants were homozygous lethal prior to adulthood, however the precise timepoint of when mutant fish died was not reported (Yoshizawa *et al*. 2011). We found that *dmrt5^sa13156^* mutants were also larval lethal (Table 1). To determine the approximate age at which *dmrt2a* and *dmrt5* mutants died, we analyzed genotypic ratios of progeny derived from heterozygous incrosses at 7 and 12dpf. For both alleles of *dmrt2a* and for *dmrt5^sa13156^*, homozygous mutants were present at 7dpf but by 12dpf homozygous mutant fish had all died (Table 2). The loss of homozygous mutants at larval stages, prior to sex-differentiation of the gonad, prevented our analysis of the function of *dmrt2a* or *dmrt5* in this process.

**Table 1.** Genotypic frequencies of adult *dmrt* mutants. Some *dmrt* mutant alleles did not follow predicted genotypic ratios, and *dmrt2a* and *dmrt5* were not adult viable; *= Chi- square test significant difference compared to expected Mendelian segregation. All fish were progeny from heterozygous crosses.

| *dmrt2a<sup>umb1</sup>* |  | *dmrt2a<sup>umb12</sup>* |  | *dmrt2b<sup>umb4</sup>* |  |
| --- | --- | --- | --- | --- | --- |
| +/+ | 23% | +/+ | *36% | +/+ | 26% |
| +/- | *77% | +/- | *64% | +/- | 49% |
| -/- | *0% | -/- | *0% | -/- | 25% |
| *dmrt2b<sup>umb6</sup>* |  | *dmrt3<sup>umb2</sup>* |  | *dmrt3<sup>umb3</sup>* |  |
| +/+ | 34% | +/+ | *38% | +/+ | 24% |
| +/- | 41% | +/- | 44% | +/- | 54% |
| -/- | 25% | -/- | *18% | -/- | 22% |
| *dmrt3<sup>umb2;umb3</sup>* |  | *dmrt5<sup>sa13156</sup>* |  |  |  |
| +/+ | *31% | +/+ | *36% |  |  |
| +/*umb2* | 22% | +/- | *64% |  |  |
| +/*umb3* | *33% | -/- | *0% |  |  |
| *umb2/umb3* | *14% |  |  |  |  |

**Table 2.**
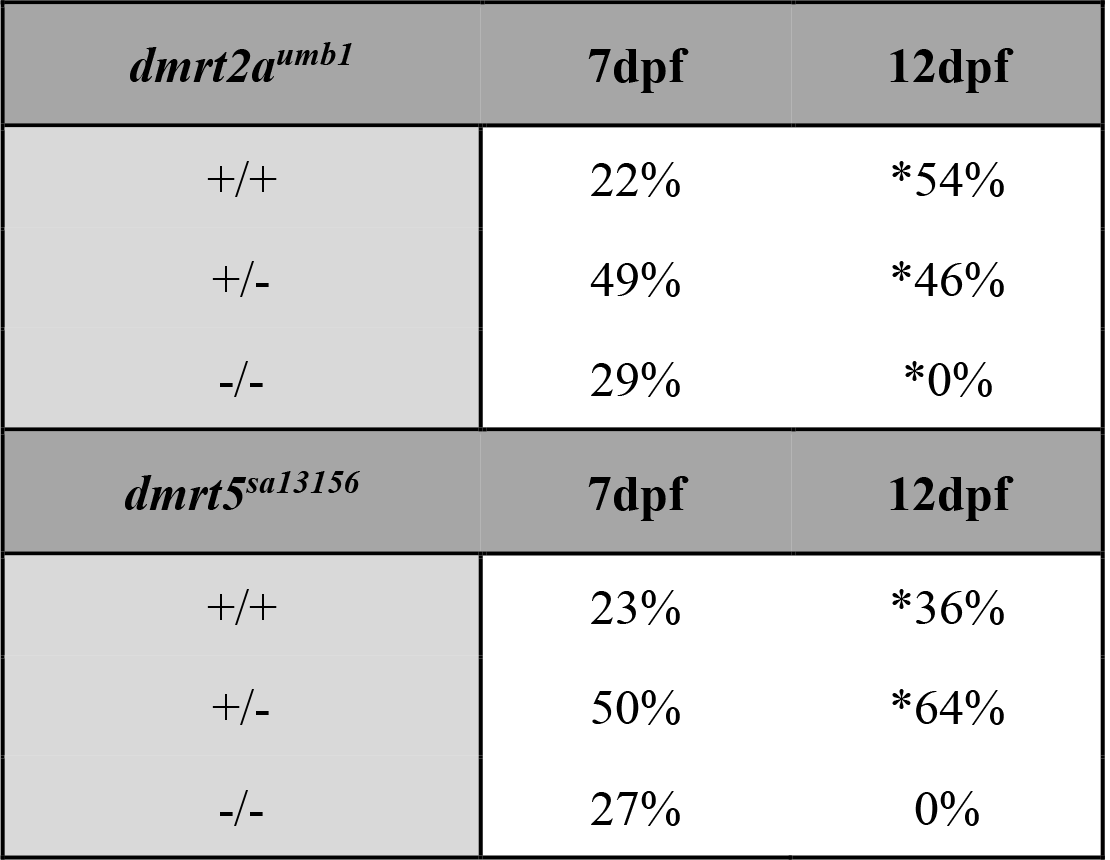
*dmrt2a* and *dmrt5* mutants are homozygous larval lethal. Mutants for *dmrt2a* and *dmrt5* died between 7 and 12dpf. *=Chi-square test significant difference compared to expected Mendelian segregation patterns. All fish were progeny from heterozygous crosses.

| *dmrt2a<sup>umb1</sup>* | 7dpf | 12dpf |
| --- | --- | --- |
| +/+ | 22% | *54% |
| +/- | 49% | *46% |
| -/- | 29% | *0% |
| *dmrt5<sup>sa13156</sup>* | 7dpf | 12dpf |
| +/+ | 23% | *36% |
| +/- | 50% | *64% |
| -/- | 27% | 0% |

### Dmrt2b and dmrt3 are dispensable for sexual development and fertility

After determining that homozygous mutants were adult viable, we asked if either *dmrt2b* or *dmrt3* have a role in sex determination or differentiation by analyzing sex ratios of adult fish. Because the strains of zebrafish used in this study do not have sex chromosomes and display variable sex ratios, sex determination defects were assayed by comparing sex ratios between wild types, heterozygotes, and mutants. Sex ratio analysis was performed on adult progeny derived from heterozygous incrosses. We found no discernable sex ratio difference between genotypes for either *dmrt2b* allele or *dmrt3^umb2^*, although sex ratios for the *dmrt3^umb3^* heterozygotes and homozygotes were slightly female biased compared to the wild types (Figure 4A,B,D,E). We next examined the sex ratios of *dmrt3^umb2/umb3^*trans- heterozygotes and found that both *dmrt3^umb2/+^* and *dmrt3^umb2/umb3^* were significantly female-biased when compared to wild-type or *dmrt3^umb3/+^* sibling populations (Figure 4G). While we did not see a significant female bias in the *dmrt3^umb2^* mutants derived from *dmrt3^umb2/+^* parents (Figure 4D), there was a slight female bias in *dmrt3^umb2/+^* fish derived from crossing *dmrt3^umb2/+^* x *dmrt3^umb3/+^* parents (Figure 4G). Given that the analysis for the trans-heterozygous mutants included ∼100 more fish than analysis of homozygous mutants, more subtle phenotypes may have been detectable. It is also possible that *dmrt3* mutations cause fish to be more sensitive to environmental influences on sex ratios, leading to variability in fish raised separately. This leads us to conclude that while loss of *dmrt2b* does not affect sex determination or differentiation, there may be subtle effects from loss of *dmrt3*.

**Figure 4.**
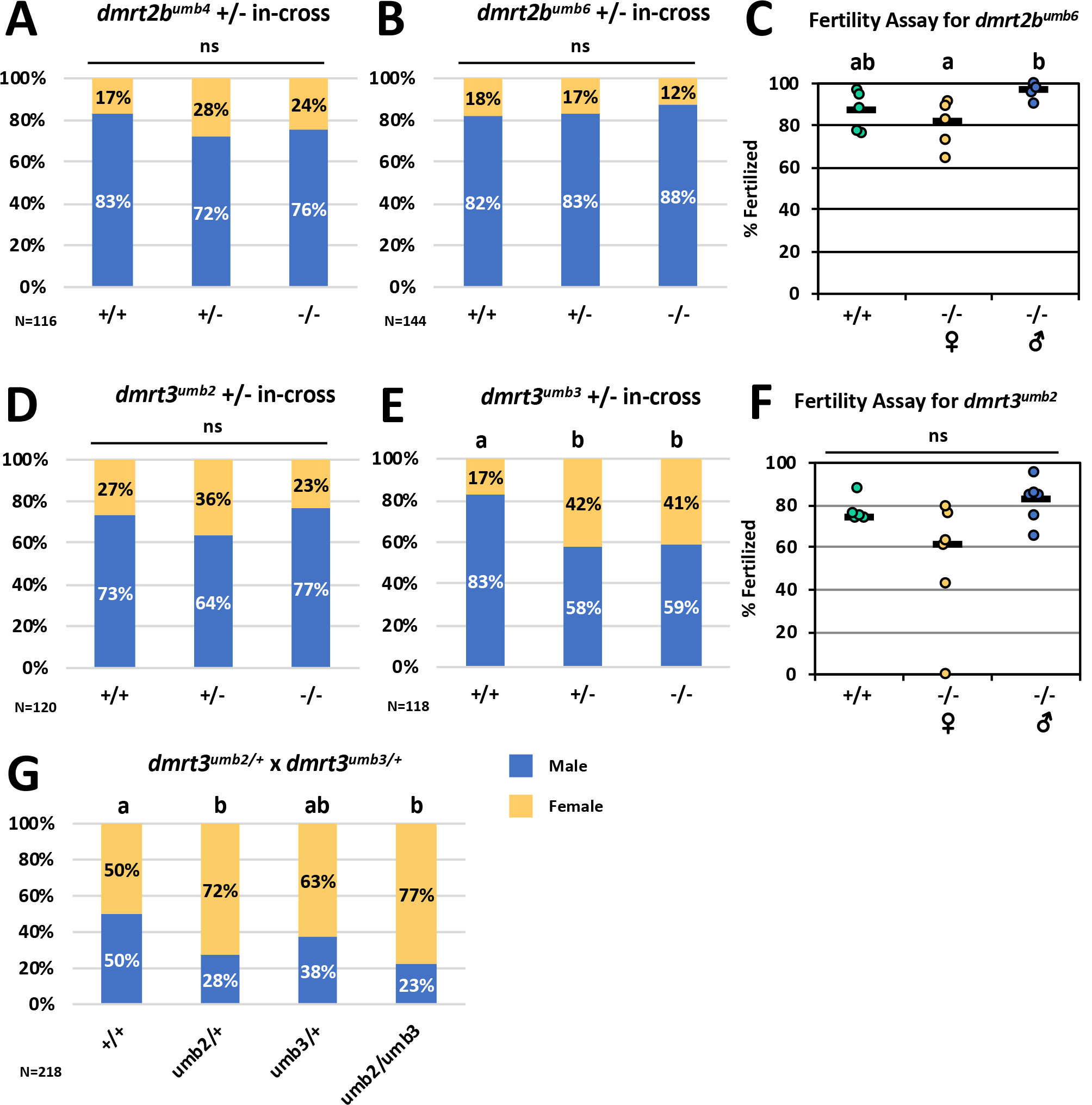
Effects of loss of *dmrt2b* or *dmrt3* on sex ratios and fertility. A-C) *dmrt2b* mutants had normal sex ratios and fertility. D-E) *dmrt3^umb2^* mutants had normal sex ratios whereas *dmrt3^umb3/+^* and *dmrt3^umb3/umb3^* were female biased compared to wild types. F) *dmrt3* mutants had normal fertility. G) Progeny from *dmrt3^umb2/+^* x *dmrt3^umb3/+^* yielded female biased sex ratios of some genotypes. Sex ratios: Fisher’s exact test for significance. Fertility: t-test for significance.

We next analyzed the fertility and gonad histology of *dmrt2b^umb6^* and *dmrt3^umb2^* mutants to ask if either gene was necessary for gonad development or function. Mutant males and females for each gene demonstrated normal fertility compared to wild types (Figure 4C,F) and produced viable progeny. The gonad histology of *dmrt2b^umb4^*, *dmrt2b^umb6^*, *dmrt3^umb2^*and *dmrt3^umb2/umb3^* mutants were histologically normal, containing all appropriate cell types and stages of germ cell development (Figure S2). We conclude that *dmrt2b* and *dmrt3* do not have major roles in zebrafish sex determination or gonad development, though *dmrt3* may have subtle effects on population sex ratios.

### Dmrt2a and dmrt3 act synergistically with dmrt1 in the zebrafish ovary

Given that *dmrt1* mutants can sometimes develop as male, we wanted to test if male development was due to redundancy with or compensation by other *dmrt* family genes. Therefore, we generated double mutants of *dmrt1* with *dmrt2a, dmrt2b* and *dmrt3* and asked if any *dmrt1* phenotypes could be modified by loss of function of one additional *dmrt* family gene.

We first analyzed *dmrt1^uc27^;dmrt2b^umb4^* double mutants to test if loss of *dmrt2b* function affected the *dmrt1* mutant phenotype. We assessed genotypic ratios of progeny resulting from *dmrt1*-/-*;dmrt2b*+/- females crossed to *dmrt1*+/-*;dmrt2b*+/- males, and determined that adult fish of all genotypic classes, including double mutants, were present in expected ratios (Table 3). We next examined the family sex ratios, asking if loss of *dmrt2b* can affect the *dmrt1* mutant female bias. We found that simultaneous loss of *dmrt1* and *dmrt2b* does not alter sex ratios (Figure 5A). To determine if loss *dmrt2b* function impacts gonad development of *dmrt1* mutants, we assayed fertility and gonad histology of *dmrt1;dmrt2b* double mutants. Double mutant females exhibited robust fertility with normal, viable progeny, and double mutant males exhibited reduced breeding and sterility, as previously described for *dmrt1* single mutants (Figure 5D) (Webster *et al*. 2017). Finally, histology for *dmrt1;dmrt2b* double mutant males and females was similar to *dmrt1*-/-, whereby females exhibited normal ovaries containing all stages of developing oocytes and males displayed aberrant testes with complete loss of normal tubule morphology and failed spermatogenesis (Figure 6 G,H). We conclude that *dmrt2b* does not act redundantly with *dmrt1* and does not modify the *dmrt1* mutant phenotype.

**Table 3.** Genotypic frequencies of double mutants. All crosses were performed by crossing *dmrt1-/-;dmrt(x)+/-* ♀ x *dmrt1+/-;dmrt(x)+/-* ♂, where (x) represents *dmrt2a, dmrt2b, or dmrt3*. Expected Mendelian segregation frequencies are shown. Because *dmrt1* is tightly linked to *dmrt2a* and *dmrt3*, these mutations do not independently assort. *Dmrt1;dmrt2b* double mutants were present at expected genotypic frequencies, but *dmrt1;dmrt2a* and *dmrt1;dmrt3* double mutants demonstrate partial lethality. *= denotes a significant deviation from expected allele segregation patterns with Chi-square test for significance.

| *dmrt1*<sup>uc27</sup><br>*dmrt2a*<sup>umb11</sup> | Expected | Observed | *dmrt1*<sup>uc27</sup><br>*dmrt2b*<sup>umb4</sup> | Expected | Observed | *dmrt1*<sup>uc27</sup><br>*dmrt3*<sup>umb14</sup> | Expected | Observed |
| --- | --- | --- | --- | --- | --- | --- | --- | --- |
| 1+/-<br>2a+/+ | 25% | 23% | 1+/-<br>2b+/+ | 13% | 11% | 1+/-<br>3+/+ | 25% | *21% |
| 1+/-<br>2a+/- | 25% | *34% | 1+/-<br>2b+/- | 25% | 20% | 1+/-<br>3+/- | 25% | *31% |
| 1-/-<br>2a+/- | 25% | *33% | 1 +/-<br>2b-/- | 13% | 14% | 1-/-<br>3+/- | 25% | *31% |
| 1-/-<br>2a-/- | 25% | *10% | 1-/-<br>2b+/- | 25% | 26% | 1-/-<br>3-/- | 25% | *17% |
|  |  |  | 1 -/-<br>2b+/+ | 13% | 13% |  |  |  |
|  |  |  | 1 -/-<br>2b-/- | 13% | 16% |  |  |  |

**Figure 5.**
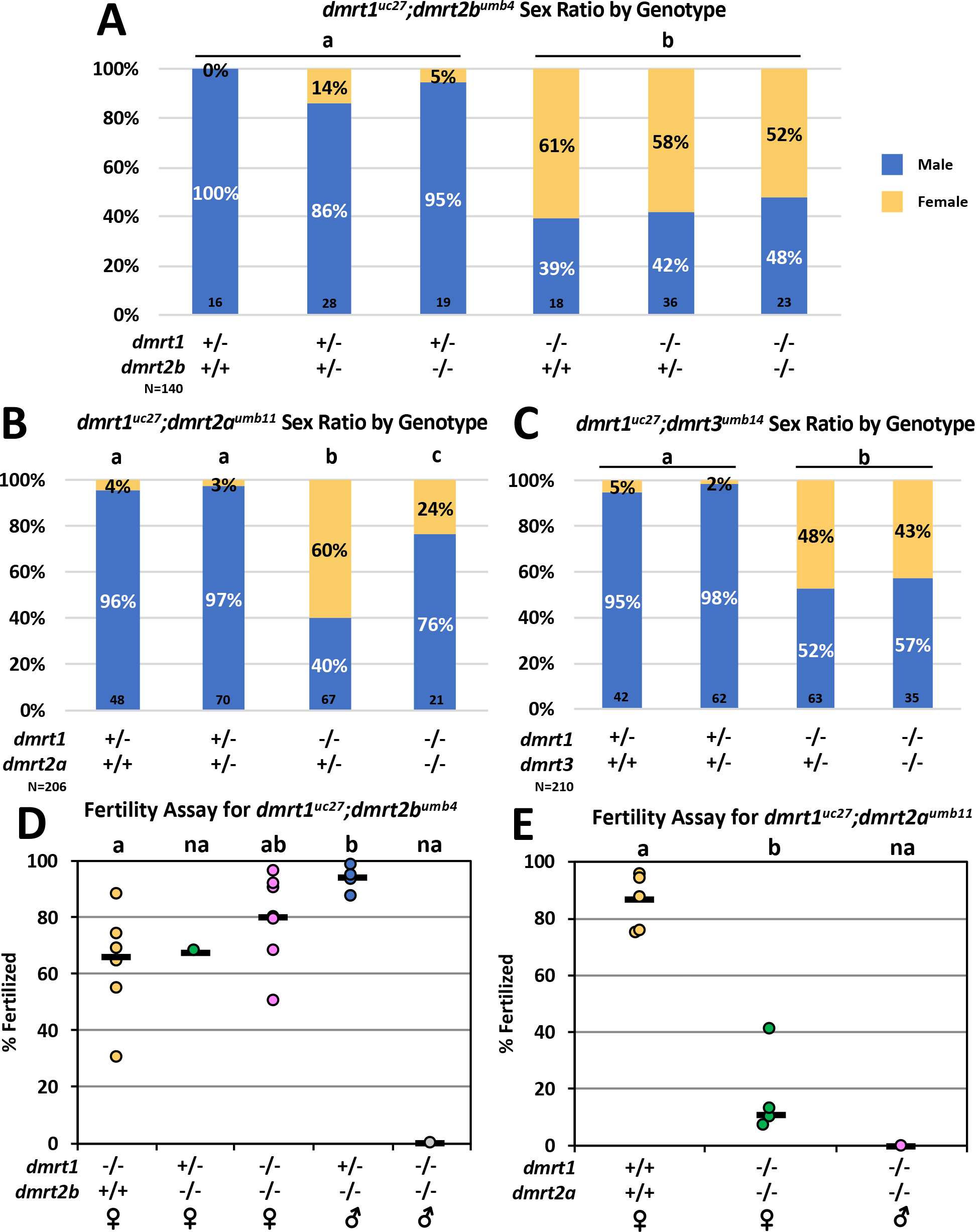
Sex ratios and fertility for *dmrt1* double mutants. A-C) Double mutants for *dmrt1* and *dmrt2b* or *dmrt3* are similar to *dmrt1* mutants, but double mutants with *dmrt2a* exhibit a partial reversal of the *dmrt1* mutant female bias. All crosses were *dmrt1-/-;dmrtx+/-* x *dmrt1+/-;dmrtx+/-* , where *dmrtx* denotes *dmrt2a, dmrt2b*, or *dmrt3.* A Fisher’s exact test was done to test significance. D) Fertility is not affected by simultaneous loss of *dmrt1* and *dmrt2b*. E) *dmrt1;dmrt2a* double mutant females are sub-fertile. D,E) Only one double mutant male induced spawning in each experiment. Significance was calculated using a T- test for fertility assays.

**Figure 6.**
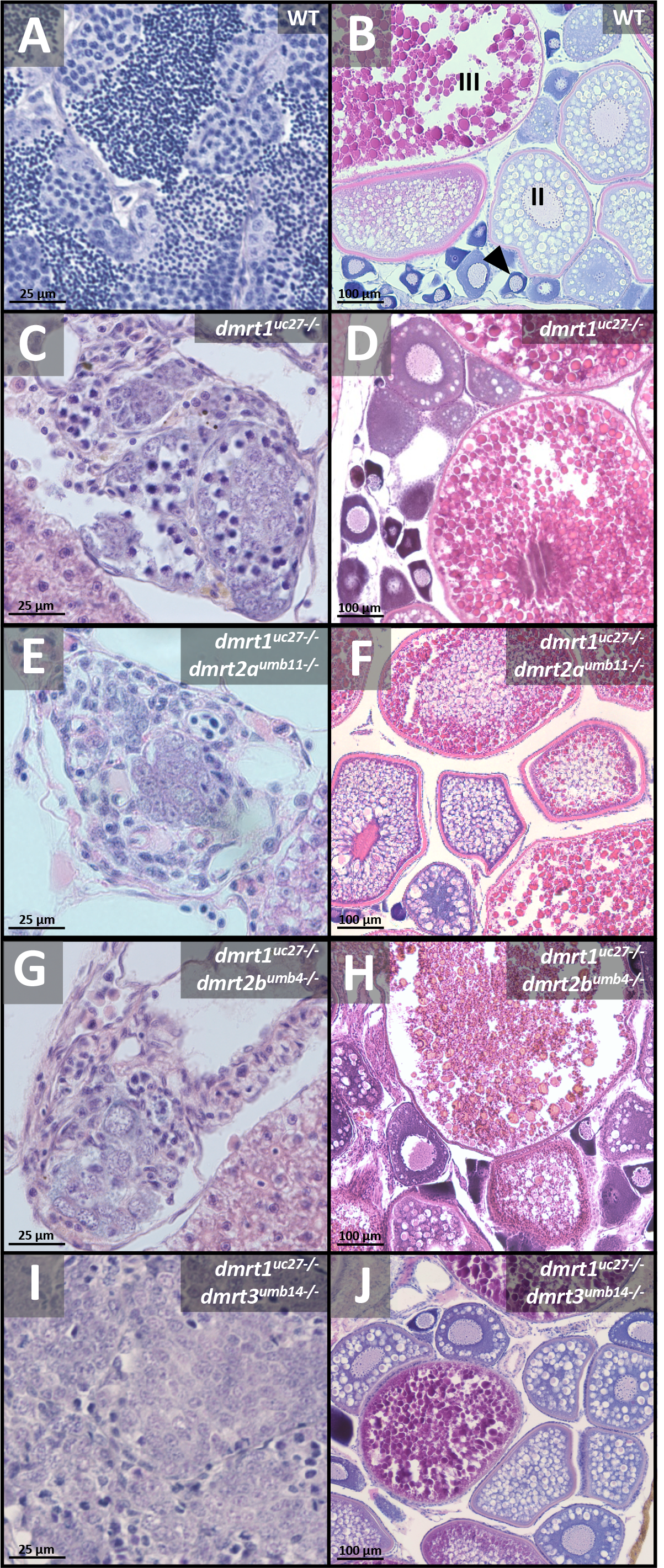
*dmrt1;dmrt2a* and *dmrt1;dmrt3* double mutants exhibit abnormal ovary morphology. H&E staining of adult ovaries and testes. Males left, females right. A-B) Wild- type testis and ovary. C-D) Dmrt1 single mutant testis and ovary. E) *dmrt1;dmrt2a* mutant testes are indistinguishable from *dmrt1-/-* males (N=3). F) *dmrt1;dmrt2a* mutant ovaries appear to have too many stage III+ oocytes and too few stage Ib (N=4). G-H) *dmrt1;dmrt2b* mutants are indistinguishable from *dmrt1-/-* (N=3). I) *dmrt1;dmrt3* testes are similar to *dmrt1* single mutants (N=3). J) *dmrt1;dmrt3* mutant ovaries appear to have too many stage II oocytes and too few stage Ib (N=4). III = stage III, II = stage, black arrow = stage Ib. Scale bars are 25μm for testes and 100μm for ovaries.

Next, we generated *dmrt2a* and *dmrt3* mutations in the *dmrt1^uc27^* mutant background to ask if mutations disrupting either of these genes could modify the *dmrt1* mutant phenotype. As *dmrt2a* mutants are homozygous lethal, we aimed to investigate if the female-biased sex ratios of *dmrt1* mutants were dominantly enhanced by having one mutant copy of *dmrt2a* (i.e., *dmrt1-/-; dmrt2a+/-).* Given that *dmrt2a* is located only 16kb from *dmrt1*, CRISPR guide RNAs targeting *dmrt2a* and Cas9-encoding mRNA were injected into *dmrt1^uc27^* fish, which yielded linked mutations to *dmrt1^uc27^*. The linked mutation *dmrt2^umb11^* is a predicted null allele with a premature stop codon at the same position as the *dmrt2a^umb1^* allele (Figure S1, Figure 3). Like *dmrt2a*, *dmrt3* is just 9kb from *dmrt1*, necessitating the generation of a novel *dmrt3* mutation in the *dmrt1^uc27^* line by CRISPR/Cas9 mutagenesis. The ensuing allele, *dmrt3^umb14^*, is also a predicted null resulting in a premature stop codon (Figure S1, Figure 3). We first analyzed the genotypic ratios of adult progeny arising from *dmrt1*-/-*;dmrt2a+/-* females crossed to *dmrt1*+/-*;dmrt2a+/-* males, and likewise for *dmrt1-/-;dmrt3+/-* females crossed with *dmrt1+/-;dmrt3+/-* males. Surprisingly, we found that *dmrt1*-/-;*dmrt2a*-/- double mutants were not absent as predicted, based on the lethality we observed for the single *dmrt2a* mutants (Tables 1,3). Instead, *dmrt1;dmrt2a* double mutants were present at 10% of the population, well-below the expected frequency of 25% (Table 3). Thus, loss of *dmrt1* function may partially rescue *dmrt2a-/-* lethality. Interestingly, *dmrt1^uc27^;dmrt3^umb14^* double mutants also exhibited some lethality, similar to that seen in *dmrt3^umb2^*mutants (Tables 1, 3). These data suggest that *dmrt1, dmrt2a*, and *dmrt3* all have roles in early development, although these roles are minor for *dmrt1* and *dmrt3*.

We next examined the family sex ratios for each line, to ask if loss of either *dmrt2a* or *dmrt3* function can modify the *dmrt1*-/- female bias. We found that *dmrt1-/-;dmrt2a+/-* fish exhibited the predicted *dmrt1* mutant female bias, relative to *dmrt1+/-;dmrt2a+/+* and *dmrt1+/-;dmrt2a+/-* siblings. By contrast, *dmrt1-/-;dmrt2a-/-* double mutants had a substantially higher proportion of males than *dmrt1-/-;dmrt2a+/-*, indicating a partial rescue of the *dmrt1-/-* male determination phenotype (Figure 5B). This suggests that *dmrt2a* functions in female sex determination. Alternatively, *dmrt2a* may be necessary for normal germ cell development, as loss of germ cells also suppresses the *dmrt1-/-* female sex bias (Webster *et al*. 2017). Simultaneous loss of *dmrt1* and *dmrt3* did not alter sex ratios relative to *dmrt1-/-;dmrt3+/-* and we conclude that there is no redundancy between *dmrt1* and *dmrt3* in determining adult sex (Figure 5C). These data point to a potential antagonistic relationship between *dmrt1* and *dmrt2a* in zebrafish sex determination, whereas *dmrt3* does not interact with *dmrt1* in this process.

We next asked if fertility of *dmrt1* mutants was affected by loss of *dmrt2a* or *dmrt3* function. Because *dmrt1-/-* males rarely breed, we could not collect robust fertility data for *dmrt1* mutant males (Figure 5D-E) (Webster *et al*. 2017). The only *dmrt1;dmrt2a* double mutant male that successfully induced spawning was infertile, as predicted for males homozygous for *dmrt1* mutations. Interestingly, double mutant females did not breed often, and displayed reduced fertility in contrast to normal fertility of *dmrt1-/-* females previously reported (Figure 5E) (Webster *et al*. 2017; Lin *et al*. 2017). Furthermore, 80% (N= two females, 1 clutch each) of the surviving progeny appeared variously abnormal and likely would not survive to maturity. Given the rarity of double mutant females in the population (2.4%), further analysis of fertility or fecundity of double mutant females was not carried out. Regardless, these data suggest that *dmrt2a* regulates ovary development. Our attempts to analyze fertility of *dmrt1;dmrt3* double mutant females were unsuccessful as these females never spawned despite numerous breeding attempts (9 pairings). These data suggest that *dmrt2a* has a role in ovary development while *dmrt3* may affect mating behavior or spawning in females.

Finally, we examined gonad histology of the *dmrt1;dmrt2a* and *dmrt1;dmrt3* double mutant gonads, to ask if morphological gonad development was affected. We found that testes from *dmrt1;dmrt2a* and *dmrt1;dmrt3* double mutant males appeared similar to the *dmrt1* mutant phenotype, indicating that loss of *dmrt2a* or *dmrt3* did not alter *dmrt1* mutant testis defects (Figure 6). By contrast, ovaries from both *dmrt1;dmrt2a* and *dmrt1;dmrt3* double mutant females were not histologically normal. Although *dmrt1*-/- ovaries appear grossly histologically normal, we found that *dmrt1-/-* fish had significantly altered proportions of oocyte stages, which was due to a decrease in stage Ib oocytes and a commensurate increase in oocytes of stages III/IV (referred to as III±; Figure 7A-D). This suggests that *dmrt1* has a function in ovary development, though it is not required for fertility (Figure 5D). The *dmrt1-/-;dmrt2a-/-* females had significantly altered proportions of oocytes compared to both wild-type and *dmrt1-/-;dmrt2a+/+* (Figure 6F, 7A). *Dmrt1-/-;dmrt2a-/-* double mutant females demonstrated a reduction of stage Ib oocytes, and a further increase in stage III± oocytes, compared to wild-type and *dmrt1-/-* fish (Figure 7C). Histology and oocyte proportions for *dmrt1;dmrt3* double mutant ovaries was also abnormal overall (Figure 6J, 7B). Analysis of the *dmrt1-/-;dmrt3-/-* double mutant females revealed a reduction of stage Ib oocytes and increased number of stage II oocytes, as compared to both wild-type and *dmrt1-/-;dmrt3+/+* fish (Figure 7D). We conclude that *dmrt1, dmrt2a* and *dmrt3* all have roles in ovary development, whereas only *dmrt1* functions in testis development.

**Figure 7.**
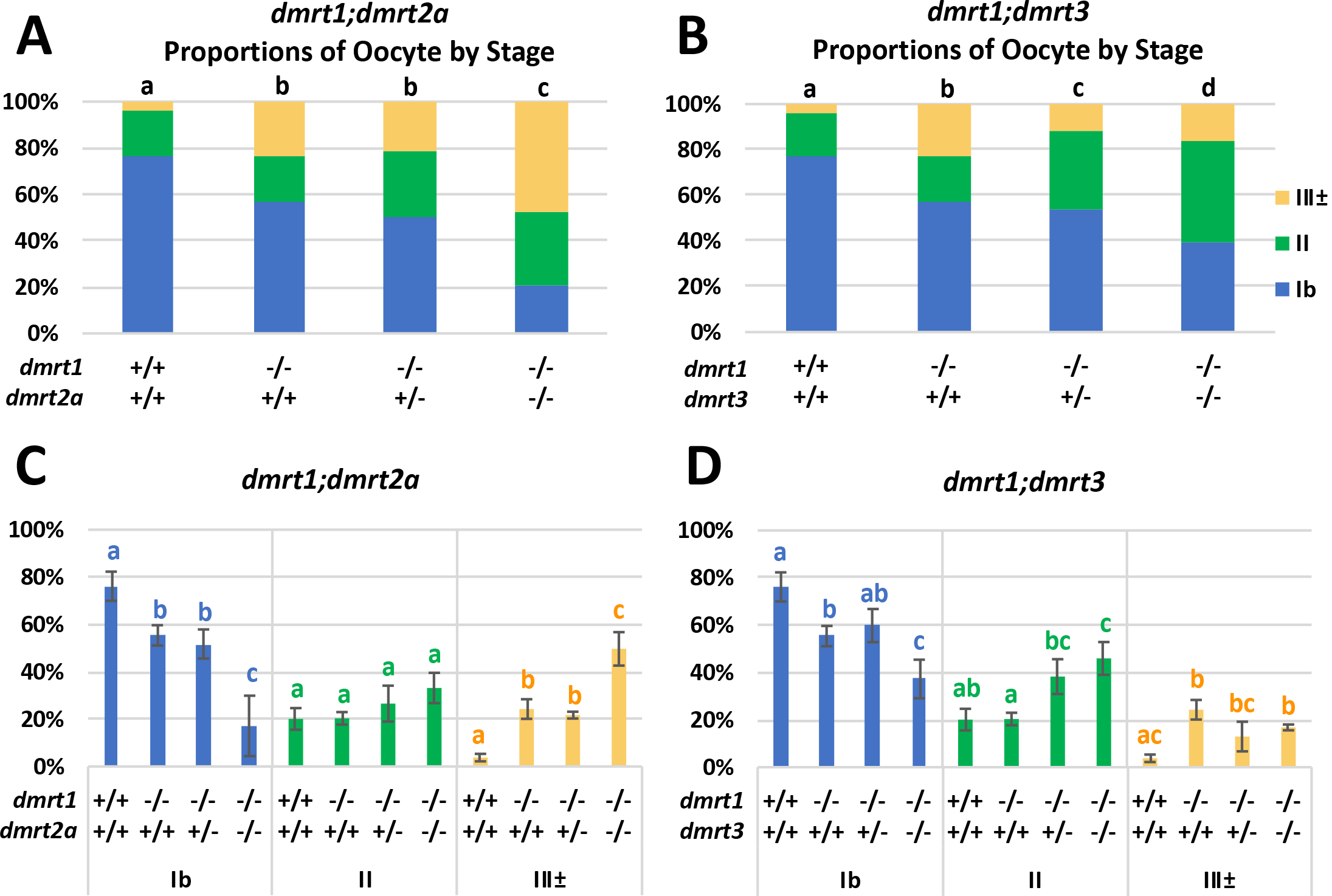
*dmrt1, dmrt2a*, and *dmrt3* function in oogenesis. Proportions of oocyte stages within ovaries from histological sections. A,B) Overall proportions of oocyte stages. Statistical significance was calculated by a chi square test. C,D) The same data is shown as in A & B, broken down for comparison of the percentage of oocytes at each stage across genotypes. A T-test was used to test significance. Wild-type and *dmrt1-/-;+/+* are the same samples used in comparison for both *dmrt1;dmrt2a* and *dmrt1;dmrt3* data sets. A) The *dmrt1* single mutants had skewed proportions of oocyte stages compared to wild types and *dmrt1;dmrt2a* had more severe differences. B) The proportions of oocytes in *dmrt1;dmrt3* double mutants is different than *dmrt1* single mutants. C-D) *dmrt1-/-;+/+* fish have reduced numbers of Ib oocytes, and increased numbers of III± oocytes compared to wild types. C) *dmrt1-/-;dmrt2a-/-* fish have reduced numbers of Ib oocytes, and increased III± oocytes as compared to wild-type and *dmrt1-/-* fish. D) *dmrt1-/-;dmrt3-/-* fish have fewer 1b and increased numbers of stage II oocytes as compared to wild-type and *dmrt1-/-;+/+* fish. III± indicates stages III and IV collectively; error bars indicate standard deviation of ratio out of each stage of oocytes out the total number of oocytes for each sample (N=3 for each genotype, except wild-type where N=2); Letters indicate significance within oocyte category.

### Dmrt1;dmrt2a and dmrt1;dmrt3 double mutants exhibit abnormal expression of key regulators of sexual development

To ask if loss of *dmrt2a* or *dmrt3* function affects expression of ovary or testis-enriched genes in the ovary, we performed qRT-PCR on *dmrt1;dmrt2a* and *dmrt1;dmrt3* double mutant ovaries in comparison to wild-type and *dmrt1-/-* ovaries. To that end, we characterized the expression of male-enriched genes *dmrt1*, *anti-müllerian hormone (amh)*, and *cytochrome P450 family 11* and *subfamily C polypeptide 1 (cyp11c1)*. We also characterized female enriched genes *cytochrome P450, family 19, subfamily A polypeptide 1a (cyp19a1a)* and *forkhead box L2a (foxl2a)*. Lastly, we examined the expression of two germ cell markers, *dead end (dnd1)*, and *DEAD (Asp-Glu-Ala-Asp) box polypeptide 4 (ddx4/vasa)*. *Amh* is a well-known regulator of sexual development and is required for normal male and female germ cell development (Lin *et al*. 2017; Yan *et al*. 2019). *Cyp11c1* is a marker for Leydig cells and is almost exclusively expressed in testes, whereas *cyp19a1a* encodes aromatase, which is required for estrogen synthesis and female sexual development (Wang and Orban 2007; Dranow *et al*. 2016; Lau *et al*. 2016; Yin *et al*. 2017; Wu *et al*. 2020; Romano *et al*. 2020). *Foxl2a* is required for maintenance of the germ cells and morphology in the mature zebrafish ovary and, together with *foxl2b*, has a role in female sex differentiation (Yang *et al*. 2017). *Dnd* is a well-known regulator of germ cells, and is required for germ cell specification and survival in zebrafish (Weidinger *et al*. 2003). Similarly, *vasa* is also a conserved regulator of germ cells, and is required for normal zebrafish germ cell differentiation (Hartung *et al*. 2014). Given the importance of this set of genes in both male and female sexual development, expression analysis of these genes could provide a better understanding of the defects observed in ovary development.

Aberrant gene expression of multiple testis-enriched genes was observed in *dmrt1;dmrt2a*, but not *dmrt1;dmrt3* double mutant ovaries. *Dmrt1* expression was reduced, as expected in *dmrt1*-/- ovaries, and added loss of *dmrt2a* led to further downregulation of *dmrt1* expression (Figure 8A). By contrast, *dmrt1;dmrt3* double mutants did not further reduce *dmrt1* expression as compared to *dmrt1-/-* ovaries. We found that *amh* expression was increased in *dmrt1-/-* ovaries relative to wild-type ovaries. This is interesting considering that *amh* expression is lost in *dmrt1-/-* testes (Webster *et al*. 2017; Lin *et al*. 2017). However, *dmrt1*-/-;*dmrt2a*-/- demonstrated a reduction of *amh* expression compared to wild-type and *dmrt1-/-* ovaries (Figure 8B). In *dmrt1-/-;dmrt3-/-, amh* expression was similar to wild-type ovaries (Figure 8B). We next analyzed *cyp11c1* expression and found that, little to no expression was detected in ovaries. We found no significant differences in *cyp11c1* expression between wild-type or double mutant ovaries, suggesting the expression of Leydig cell genes are not affected by loss of *dmrt2a* or *dmrt3* (Figure 8C). Overall, mis-expression of *dmrt1* and *amh* in *dmrt1-/-* ovaries was affected by loss of *dmrt2a* function but not *dmrt3*.

**Figure 8.**
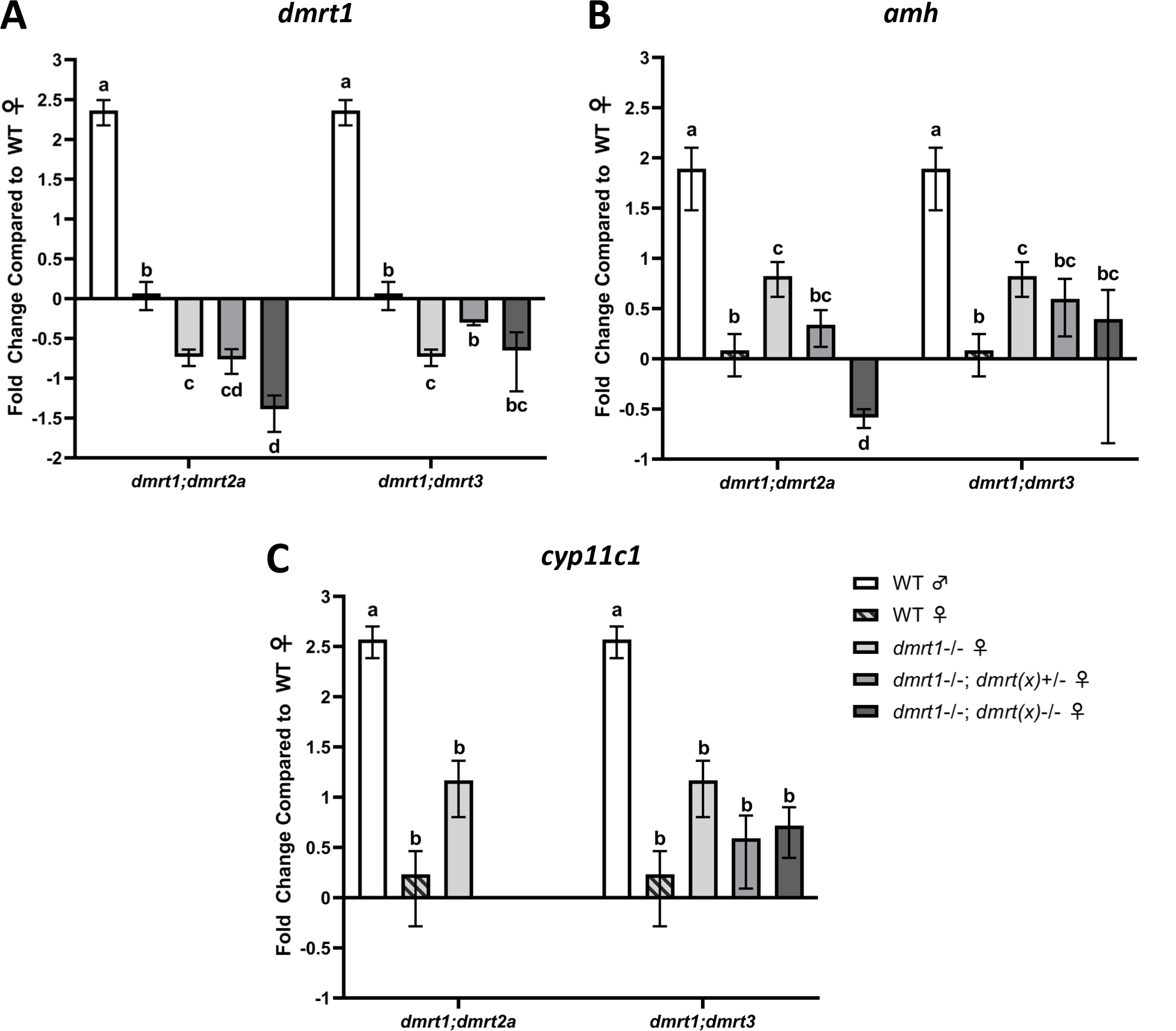
Genes with testes enriched expression mis-expressed in *dmrt1;dmrt2a* but not *dmrt1;dmrt3* ovaries. qRT-PCR for A) *dmrt1*, B), *amh*, and C) cyp11c1 gene expression in adult ovaries. *dmrt(x)* indicates either *dmrt2a* or *dmrt3*. Wild-type testes and ovaries, and *dmrt1-/-* ovaries are the same samples for both *dmrt1;dmrt2a* and *dmrt1;dmrt3*, and included in both for comparison purposes. All other fish within *dmrt1;dmrt2a* or *dmrt1;dmrt3* groups are siblings. Letters specify significance.

We next examined expression of ovary-enriched genes in *dmrt1-/-*, and *dmrt1-/-* double mutant for either *dmrt2a* or *dmrt3.* We found that while loss of *dmrt1* did not affect *cyp19a1a* expression when compared with wild-type ovaries, *cyp19a1a* expression was reduced by partial (*dmrt1-/-;dmrt2a+/-)* or complete (*dmrt1-/-;dmrt2a-/-)* loss of *dmrt2a* function (Figure 9A). In *dmrt1-/-;dmrt3-/-* ovaries *cyp19a1a* expression was similar to *dmrt1-/-* (Figure 9A). *Foxl2a* expression was decreased in *dmrt1-/-;dmrt2a+/-* and *dmrt1-/-;dmrt2a-/-* ovaries compared to *dmrt1*-/- ovaries (Figure 9B). Similarly, *dmrt1-/-;dmrt3+/-* and *dmrt1-/-;dmrt3-/-* ovaries had reduced *foxl2a* expression compared to *dmrt1-/-*. In summary, expression of *cyp19a1a* and *foxl2a* was affected by simultaneous loss of *dmrt1* and *dmrt2a* or *dmrt3* function compared to *dmrt1* single mutants indicating that these *dmrt* genes may act cooperatively in ovary development.

**Figure 9.**
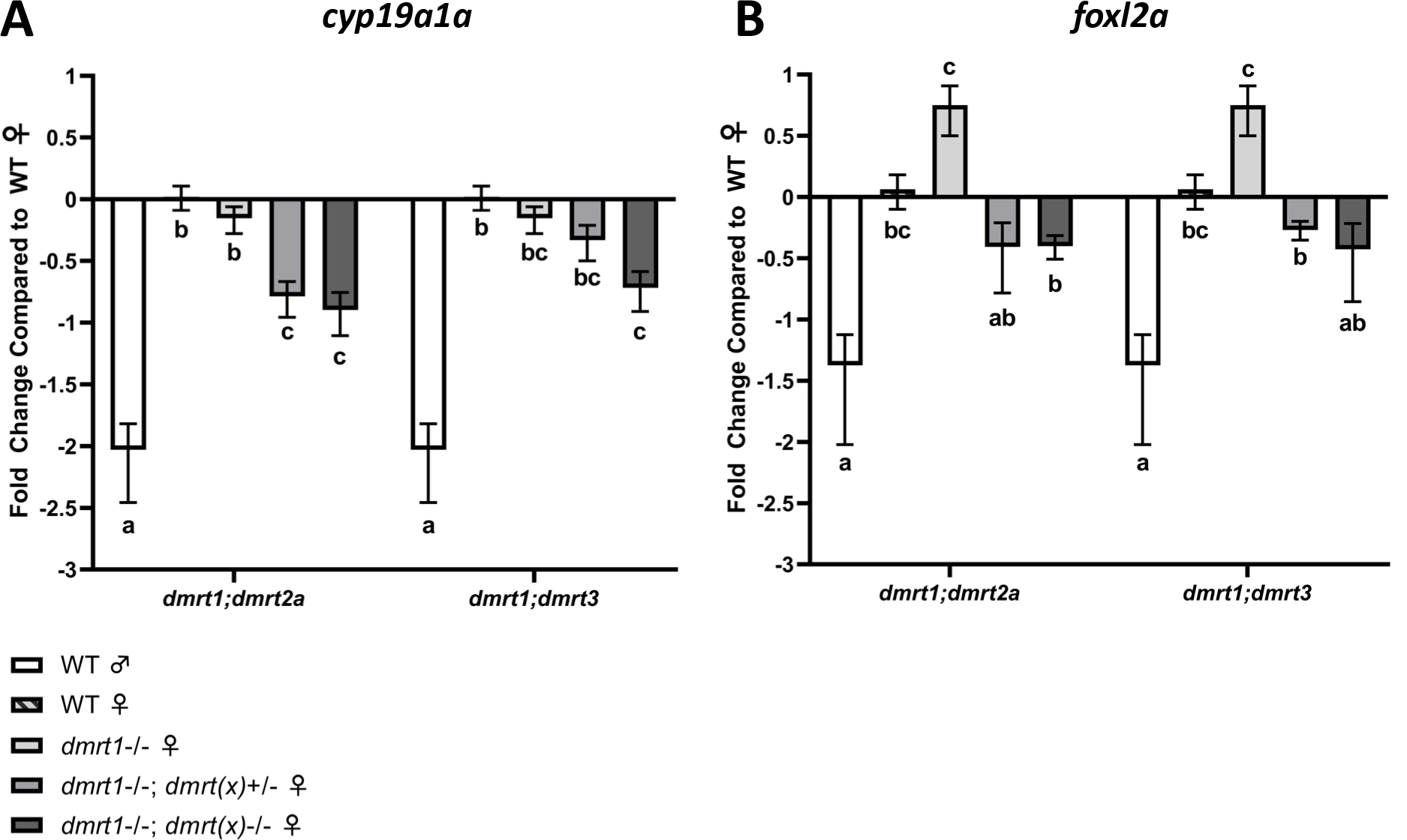
Genes commonly involved in female sexual development are mis-expressed in *dmrt1;dmrt2a* and *dmrt1;dmrt3* ovaries. qRT-PCR A) *cyp19a1a* and B) *foxl2a* gene expression in ovaries from *dmrt1* single mutants and *dmrt1;dmrt2a* and *dmrt1;dmrt3* double mutants. *dmrt(x)* indicates either *dmrt2a* or *dmrt3*. Wild-type testes and ovaries, and *dmrt1-/-* ovaries are the same samples for all experiments and included in duplicate for comparison purposes. All other fish within *dmrt1;dmrt2a* or *dmrt1;dmrt3* groups are siblings. Letters specify significance within *dmrt1;dmrt2a* and I sample groups but not between them.

We also investigated the expression of two germ cell-expressed genes, *dnd1* and *vasa. Dmrt1-/-;dmrt2a-/-* and *dmrt1-/-;dmrt3-/-* ovaries had decreased expression of *dnd1* relative to wild-type and *dmrt1-/-* ovaries (Figure 10A). In contrast to what was observed for *dnd1* expression, *vasa* expression was increased in *dmrt1-/-* ovaries compared to wild type. We found that *vasa* expression was reduced in *dmrt1-/-;dmrt2a-/-* ovaries, similar to *dnd1* (Figure 10B). By contrast, *dmrt1-/-;dmrt3-/-* ovaries had similar levels of *vasa* expression to that of *dmrt1-/-* (Figure 10B). Differences in how these mutations affect expression of *dnd1* and *vasa* in ovaries could reflect differences in the oocyte stages that they are expressed in, however such expression data has only been reported for *vasa* in adult zebrafish ovaries (Howley and Ho 2000; Kosaka *et al*. 2007). Collectively, these data suggest that *dmrt2a* and *dmrt3* function in ovary development, particularly in proper differentiation of female germ cells.

**Figure 10.**
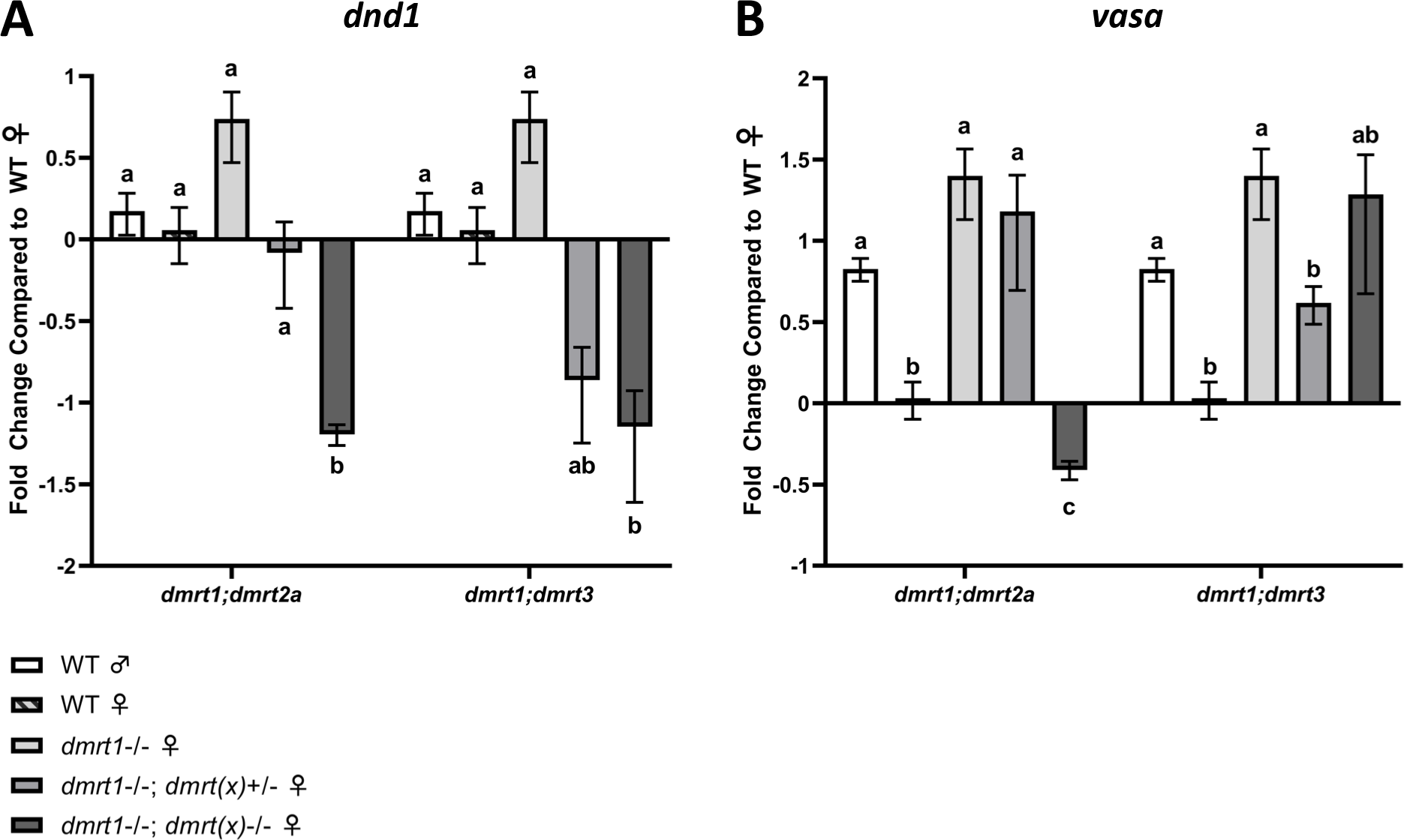
Genes involved in germ cell development are mis-expressed in *dmrt1;dmrt2a* and *dmrt1;dmrt3* ovaries. qRT-PCR for A) *dnd* and B) *vasa* expression, dmrt(x) indicates either dmrt2a or dmrt3. Wild-type testes and ovaris, and *dmrt1-/-* ovaries are the same samples for both *dmrt1;dmrt2a* and *dmrt1;dmrt3* in each chart, and are included in both for comparison purposes. All other ovaries within *dmrt1;dmrt2a* or *dmrt1;dmrt3* groups are from siblings. Letters specify significance within *dmrt1;dmrt2a* and *dmrt1;dmrt3* groups but not between them.

## DISCUSSION

The aim of this study was to assess the function of members of the zebrafish *dmrt* gene family in gonadal sexual development and their possible redundancy with *dmrt1*. We found that *dmrt2b* appears dispensable for sexual development, as *dmrt2b* mutants had unaltered sex ratios with normal fertility and gonad histology. Although *dmrt2a* single mutants were homozygous lethal, a small percentage of mutants survived when double mutant with *dmrt1.* Loss of *dmrt2a* function partially rescued the female bias of *dmrt1* mutants, suggesting that *dmrt2a* acts antagonistically to *dmrt1* in female sex differentiation. We identified previously unknown roles of *dmrt1* in ovary development, with mutant ovaries having an increase in stage Ib and decrease in stages III± follicles, although fertility was normal. *Dmrt1;dmrt2a* double mutant females displayed an enhanced reduction of stage Ib and an overabundance of stage III± oocytes, and impaired fertility, compared to wild-type and *dmrt1* single mutants. Similarly, *dmrt1;dmrt3* double mutants demonstrated a reduction of stage Ib oocytes, an increase in stage II oocytes, and failure to breed. Furthermore, we showed that double mutant ovaries for *dmrt1;dmrt2a* and *dmrt1;dmrt3* both exhibited reduced expression of female-enriched genes, and aberrant expression of germ cell markers. Taken together, these data place *dmrt2a* function in opposition to *dmrt1* to promote female sex determination, while both *dmrt2a* and *dmrt3* may act cooperatively with *dmrt1* in oocyte development (Figure 11).

**Figure 11.**
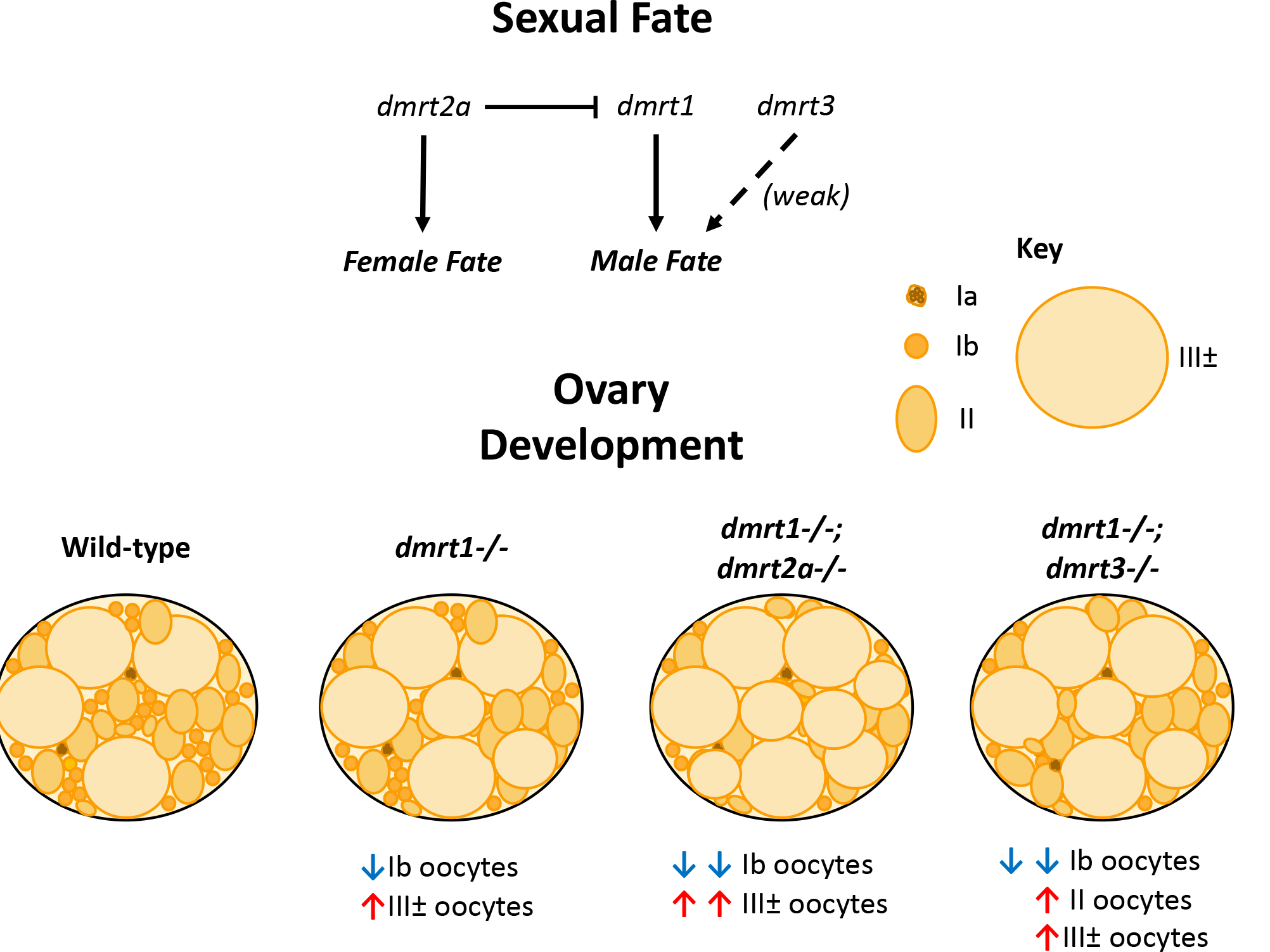
Zebrafish *dmrt* genes have antagonistic and synergistic functions in sexual development. Top: Similar to *dmrt1*, *dmrt3* is involved in specifying male fate, though it is not required for male development. Conversely, *dmrt2a* is antagonistic to *dmrt1* to promote female fate. Bottom: Mutant analysis revealed roles of *dmrt1, dmrt2a*, and *dmrt3* in oogenesis. *Dmrt1* loss of function results in increased stage III± oocytes and reduction in Ib oocytes. This is exacerbated by loss of *dmrt2a*, where the Ib population is even further reduced, and the III± population is further increased. Loss of *dmrt1;dmrt3* double mutants also results in further loss of Ib oocytes and an increase in II rather than III± oocytes.

### Zebrafish dmrt2a, dmrt2b, and dmrt3 have different developmental roles

In mice, *Dmrt2* has a well described roles in embryogenesis, particularly somite development (Seo *et al*. 2006). Zebrafish have two *dmrt2* paralogous, which arose prior to the teleost genome duplication (Zhou *et al*. 2008; Johnsen and Andersen 2012). We found that *dmrt2a* mutants were larval lethal whereas *dmrt2b* mutants were adult viable and fertile. The *dmrt2a* mutations described here caused lethality between 5 and 7 dpf, indicating that it has an essential role in early development, although we did not investigate the cause. As these larvae were able to swim after 5 dpf and appeared morphologically normal, there are likely no severe early somite defects. Surprisingly, *dmrt2a^umb11^* mutants were only sub-viable as double mutants with *dmrt1*. This could be due to numerous reasons, including: 1) the *dmrt2a^umb11^*allele generated in the *dmrt1* mutant line could be functioning as a hypomorph, rather than a true null allele; 2) there could be rescue effects caused by the loss of *dmrt1*; 3) other genetic differences between the *dmrt1* mutant line and our AB wild-type line confer partial viability of *dmrt1a* mutants. Given that multiple single *dmrt2a* alleles had complete lethality (Table 1), it seems unlikely that *dmrt2a^umb11^* is a less severe mutation compared to the other two alleles, considering that the premature stop codon produced in the *dmrt2a^umb11^* allele is identical to that of *dmrt2a^umb1^* (Figure S1, Figure 3). It is probable that the rescue is due to either loss of *dmrt1* function or another modifier in the background of the *dmrt1* mutant line.

Zebrafish *dmrt2a* and *dmrt2b* mutants and morphants have been previously reported, some of which suggest roles in embryonic development. In contrast to the *dmrt2a-/-* lethality we report here, prior work suggested that zebrafish *dmrt2a* mutants were normal and viable, with only a small number of embryos exhibiting left-right patterning defects (Pinto *et al*. 2018). Morpholino knockdowns suggested roles in left-right patterning, synchronization of the left and right somite clock, and development of fast and slow muscle (Saúde *et al*. 2005; Lu *et al*. 2017; Pinto *et al*. 2018). The discrepancies between the two viable mutations reported by Pinto et al. and our two lethal mutations and one sub-viable mutation are not easily explained – the viable mutations disrupt the translational start, whereas our mutations are about 80bp downstream of the start codon. The *dmrt2b* mutants reported in this study were adult viable and exhibited no effects on sexual fate, fertility, or gonad development. Previously, knockdowns with morpholinos targeting *dmrt2b* suggested that it is required for somitogenesis, muscle, and heart development (Liu *et al*. 2009). A mutation disrupting *dmrt2b* exhibited defects in pharyngeal pouch, cranial neural crest, and facial cartilage development, while a second *dmrt2b* mutation had no apparent defects, in agreement with our two alleles (Li *et al*. 2018; Pinto *et al*. 2018). All four alleles (2 reported here and 2 previously described) disrupt the gene in a similar region within 36 base pairs of each other, making it unclear why some of these mutations have dramatically different phenotypic consequences. Additional mutations that delete the entire locus of each of these genes or block production of a transcript will be needed to discover the true null phenotypes.

We found *dmrt3* mutants to be normal and fertile with a moderate female bias detected in some experiments (Figure 4). The viable phenotype we report agrees with recently reported mutants, which are viable with larval locomotion and juvenile swim performance defects (Del Pozo *et al*. 2020). Similar to our results in zebrafish, loss of *Dmrt3* does not appear to alter viability or fertility in mice, although a link between *DMRT3*, *2’-5’- Oligoadenylate Synthetase 3* (*OAS3*), *Estrogen Receptor 1* (*ESR1*), and human sexual differentiation has been reported (Inui *et al*. 2017; Tsai *et al*. 2020). *OAS3* does not exist outside of mammals, thus this particular role for human *DMRT3* may have evolved independently, or may be performed by alternatives to *OAS3* outside of mammals. The female-biased sex ratios we found in some, but not all families suggests that the role of *dmrt3* in zebrafish sexual development is subtle and may be sensitive to environmental or genetic modifiers. Given these data in humans and zebrafish, subtle roles for *dmrt3* in sexual development may be conserved in vertebrates.

### Dmrt1, dmrt2a and dmrt3 function in oocyte development

Infertility is a global problem significantly on the rise, with up to 19% of couples reporting problems with infertility in the US (female partner age 15-49 years old) (Centers for Disease Control and Prevention). Click or tap here to enter text.However, causes of infertility are not well understood, with 30-40% of couples diagnosed with “unexplained infertility” (Collins and Crosignani 1992). Thus, the investigation of subtle and redundant effects between genes is paramount, as problems in fertility are more likely to be a spectrum, existing in a delicate system of checks and balances, rather than isolated obstacles. Underscoring this, through double mutant analysis of *dmrt1;dmrt2a* and *dmrt1*;*dmrt3* we identified roles in oocyte differentiation for these genes.

*Dmrt1* is known to be expressed in Ib oocytes, although *dmrt1* mutant females have normal fertility and fecundity (Webster *et al*. 2017; Lin *et al*. 2017). We found that *dmrt1* mutants had decreased numbers of Ib oocytes and an increase in the number of stage III± oocytes compared to wild types (Figure 7B-E). In addition, *amh* expression was increased relative to wild-type ovaries, which is expressed in follicle cells of stage Ib, II, and III oocytes (Rodríguez-Marí *et al*. 2005). By contrast *cyp19a1a* and *foxl2a*, which are expressed in similar staged follicle cells to *amh*, were not affected (Rodríguez-Marí *et al*. 2005; Dranow *et al*. 2016; Yang *et al*. 2017). *Amh* expression is also disrupted in *dmrt1-/-* testes, although it has reduced expression in mutant testes, suggesting that *dmrt1* is important for proper *amh* expression in both sexes (Webster *et al*. 2017; Lin *et al*. 2017). Because *amh* and *dmrt1* are not expressed in the same cell types in ovaries, this relationship is indirect. Here we report that *dmrt1* has a heretofore uncharacterized function in the ovary, although is not necessary for robust female fertility.

*Dmrt1* mutant ovaries exhibited more severe defects when double mutant with either *dmrt2a* or *dmrt3.* Double mutants had increased altered proportions of oocyte stages than *dmrt1-/-*, with both having further reductions in stage 1b follicles compared to *dmrt1-/-*, while only *dmrt2a* loss further enhanced the increase of stage III follicles seen in *dmrt1-/-* ovaries. Expression of genes found in follicle cells were generally down-regulated in double mutants compared to *dmrt-/-. Amh, foxl2a*, and *cyp19a1a* expression were all reduced in *dmrt1;dmrt2a* double mutants compared to *dmrt1* single mutants, whereas only *foxl2a* was downregulated in *dmrt1;dmrt3* double mutants. In addition, *dmrt1;dmrt2a* double mutant females had reduced fertility and abnormal progeny, in contrast to normal the fertility and fecundity of *dmrt1* mutant females. Interestingly, double mutants for *dmrt1;dmrt2a* were not rigorous breeders, and we were unsuccessful in our attempts to breed *dmrt1;dmrt3* females entirely. This suggests that simultaneous loss of *dmrt1;dmrt2a* or *dmrt1;dmrt3* results in oocyte differentiation defects and may also affect female breeding success. Taken together, these results demonstrate that not only does *dmrt1* function in the zebrafish ovary, it also functions synergistically in that capacity with *dmrt2a* and *dmrt3*. While not widely investigated, *dmrt2* expression has been detected in the ovary of several other species, for example (Zhou *et al*. 2008; Shi *et al*. 2014; Zhu *et al*. 2019; Zhang *et al*. 2022), suggesting that the function of *dmrt2a* in the ovary may be conserved in other organisms.

Multiple *dmrt* genes, such as *Dmrt1, Dmrt6* and *Dmrt7*, have reported roles in the gonad and often are associated with defects in the testes (Kawamata and Nishimori 2006; Kim *et al*. 2007b; Zhang *et al*. 2014a, 2014b; Li *et al*. 2020). This conservation of *dmrt* gene function in male development is maintained even beyond vertebrates. In *Drosophila*, the *dmrt11E* gene is expressed in the somatic cells of the testis and is required for spermatogenesis and male fertility (Yu *et al*. 2015). By contrast, *doublesex (dsx)*, from which the *dmrt* family partially derives its name, is necessary for sex determination but not directly involved in gametogenesis (Casper and Van Doren 2006). In arthropods, which lack *dmrt2, dmrt11E* is a closely related orthologue, suggesting that functions for this gene cluster in sexual development are potentially ancestral (Mawaribuchi *et al*. 2019).

Previous studies have also revealed roles for *dmrt* genes in the ovary. Loss of *Dmrt4* has been shown to cause polyovular follicles in mice, though it does not alter their fertility, and xenograft experiments have revealed a role for *DMRT5* in human oogonia differentiation (Balciuniene *et al*. 2006; Poulain *et al*. 2014). Interestingly, this study also implicated involvement for *DMRT6* and *DMRT7* in the ovary, as the expression of both were impaired when *DMRT5* was inhibited in the fetal ovary (Poulain *et al*. 2014). Finally, while *dmrt11E* is necessary for spermatogenesis in *Drosophila*, in the domesticated silkworm (Bombyx mori), *dmrt11E* is instead required for normal oogenesis and female fertility (Kasahara *et al*. 2021). Given that *dmrt11E* is an orthologue to *dmrt2*, this suggests that germ cell development may be a conserved ancestral function for these genes. Our work on zebrafish shed light on the subtle and redundant effects of genes involved in ovary development and will be important to better characterize, and eventually treat, various causes of infertility.

### Conclusions

Here we have presented the first evidence for *dmrt* gene function in the zebrafish ovary. We have shown that, unlike *dmrt1*, single mutants for *dmrt2b* and *dmrt3* are generally not disruptive to normal zebrafish sexual development. However, we found that simultaneous loss of *dmrt1* and *dmrt2a* results in partial reversal of the *dmrt1*-/- sex determination phenotype. Furthermore, double mutants for *dmrt1* and *dmrt2a* or *dmrt3* exhibit more severe defects in oocyte development than *dmrt1* single mutants, as well as subfertility and failure to breed, respectively. Taken together, these results show that *dmrt1* acts synergistically with *dmrt2a* and *dmrt3* in female germ cell development, while *dmrt1* and *dmrt2a* act antagonistically in determining zebrafish sexual fate.

## ACKNOWLEDGEMENTS

We would like to thank all the members, past and present, of the Siegfried Lab at UMB for their support and helpful discussions. Carlee Dawson and Courtney Aslan-Rollins both provided technical support for this study. We also thank the Zebrafish International Resource Center (ZIRC) for providing the *dmrt5^sa13156^* line.

## FIGURE LEGENDS

**Figure S1.** Genomic sequence showing *dmrt* gene mutations, with 10bp flanking sequence included for reference. grey = flanking genomic sequence, black = affected wild-type genomic sequence; blue = insertion, green = missense base change, red dash = deleted base

**Figure S2. *Dmrt2b* and *dmrt3* mutants have normal gonad morphology.** Ovaries left, testes right. H&E staining of adult ovaries and testes; N=3 for genotypes. Scale bars are 25μm for testes and 100μm for ovaries.

